# Isolation of Chemically Cyclized Peptide Binders Using Yeast Surface Display

**DOI:** 10.1101/2020.04.16.044438

**Authors:** Kaitlyn Bacon, Abigail Blain, Matthew Burroughs, Nikki McArthur, Balaji M. Rao, Stefano Menegatti

## Abstract

Cyclic peptides with engineered protein-binding activity have gained increasing attention for use in therapeutic and biotechnology applications. We describe the efficient isolation and characterization of cyclic peptide binders from genetically encoded combinatorial libraries using yeast surface display. Here, peptide cyclization is achieved by disuccinimidyl glutarate-mediated crosslinking of amine groups within a linear peptide sequence that is expressed as a yeast cell surface fusion. Using this approach, we first screened a library of cyclic heptapeptides by magnetic selection and fluorescence activated cell sorting (FACS), to isolate binders for a model target (lysozyme) with low micromolar binding affinity (K_D_ ~ 1.2 - 3.7 µM). The isolated peptides bound lysozyme selectively, and only when cyclized. Importantly, we showed that yeast surface displayed cyclic peptides could be used to efficiently obtain quantitative estimates of binding affinity, without chemical synthesis of the selected peptides. Subsequently, to demonstrate broader applicability of our approach, we isolated cyclic heptapeptides that bind human interleukin-17 (IL-17) using yeast-displayed IL-17 as a target for magnetic selection, followed by FACS using recombinant IL-17. Molecular docking simulations and follow-up experimental analyses identified a candidate cyclic peptide that binds IL-17 in its receptor binding region with moderate affinity (K_D_ ~ 300 nM). Taken together, our results show that yeast surface display can be used to efficiently isolate and characterize cyclic peptides generated by chemical modification from combinatorial libraries.

## Introduction

Cyclic peptides have emerged as a promising class of affinity ligands for use in basic research as well as diagnostic and therapeutic applications due to their favorable properties such as higher binding affinity and stability relative to linear peptides^1^. In particular, they are ideal candidates to inhibit protein-protein interactions as their size and modularity allows them to closely mimic the principal attributes of these normally flat, featureless interaction surfaces. Other attractive features of cyclic peptides include scalable and affordable synthesis, metabolic stability, and biocompatibility. Additionally, they can be easily modified with labels (*e.g.*,fluorescence or radioactive probes)^2^ or conjugated to other biomolecules^3^ to endow further biochemical functionalities.

Cyclic peptides are commonly isolated from combinatorial libraries that are either chemically synthesized or genetically encoded. Among chemical synthesis-based approaches, one-bead-one-peptide libraries are frequently employed^4^; these libraries are a collection of beads, each displaying multiple copies of a unique peptide sequence that can be sequenced by Edman degradation or mass spectrometry. Commonly used methods for generating genetically-encoded peptide libraries include mRNA display^5^, ribosomal display^6^, and phage-display^7^. Multiple strategies have been developed for the cyclization of displayed peptides, both chemical^8^ and enzymatic^9^, although disulfide bond formation remains the most widely employed^10^. Disulfide-containing cyclic peptides, however, are unlikely to maintain a stable cyclic structure *in vivo* due to the reducing environment of the cytosol. Alternative routes to stable peptide cyclization involve the formation of thioether and amide bonds via chemical crosslinking, which have been demonstrated with mRNA display^11^ and phage-display libraries^8^.

Chemically synthesized and genetically encoded peptide libraries are typically screened using a “panning” method, in which the library is incubated with a surface where the target protein is immobilized, and clones that bind the surface are isolated. However, panning-based selections do not provide an efficient means to discriminate high-affinity binders from low affinity binders^12^. Further, panning methods typically identify a pool of peptides that subsequently need to be chemically synthesized to characterize biochemical properties, like binding affinity. To overcome these limitations, we describe the use of yeast surface display for the construction of chemically cyclized peptide libraries that can be subjected to fluorescence activated cell sorting (FACS) to efficiently isolate the highest affinity library clones. Only a few instances are reported of using yeast display to express cyclic peptides, most of which are limited to disulfide-constrained peptides, like knottins^13^. However, to our knowledge, chemical crosslinkers have not been commonly used for the cyclization of yeast-displayed peptides, despite their widespread use for cyclizing mRNA- and phage-display peptide libraries.

In our approach, a linear peptide precursor is displayed on the yeast cell surface as an N-terminal fusion to the Aga2p subunit of the yeast mating protein α-agglutinin. The Aga2p subunit forms a disulfide bond with the Aga1p subunit of α-agglutinin, thereby tethering the peptide to the yeast cell surface. Peptide cyclization occurs by crosslinking the N-terminal amine and the ɛ-amine of a lysine residue (Figure 1). Amino acid residues between an N-terminal methionine and a C-terminal lysine are randomized to generate a combinatorial library. It is important to note that a prepro secretory signal sequence precedes the linear peptide precursor. A portion of this sequence directs the nascent polypeptide through the secretion pathway involving the endoplasmic reticulum and the Golgi apparatus^14^. Before leaving the Golgi, the secretory sequence is cleaved between its C-terminal lysine and arginine residues^15^, resulting in an arginine residue at the N-terminus of the displayed peptide. Thus, the displayed peptides are crosslinked between the α-amine of arginine and the ɛ-amine of a lysine located at the C-terminus of the peptide sequence. Other components of the displayed peptide fusion include a HA epitope tag to characterize the expression of the fusion; no lysine residues are present in this tag. Whilst Aga2p includes three lysine residues, DSG-mediated crosslinking between these lysine residues and amine groups within the linear precursor peptide is unlikely due to the large spatial separation between these moieties.

**Figure 1.**
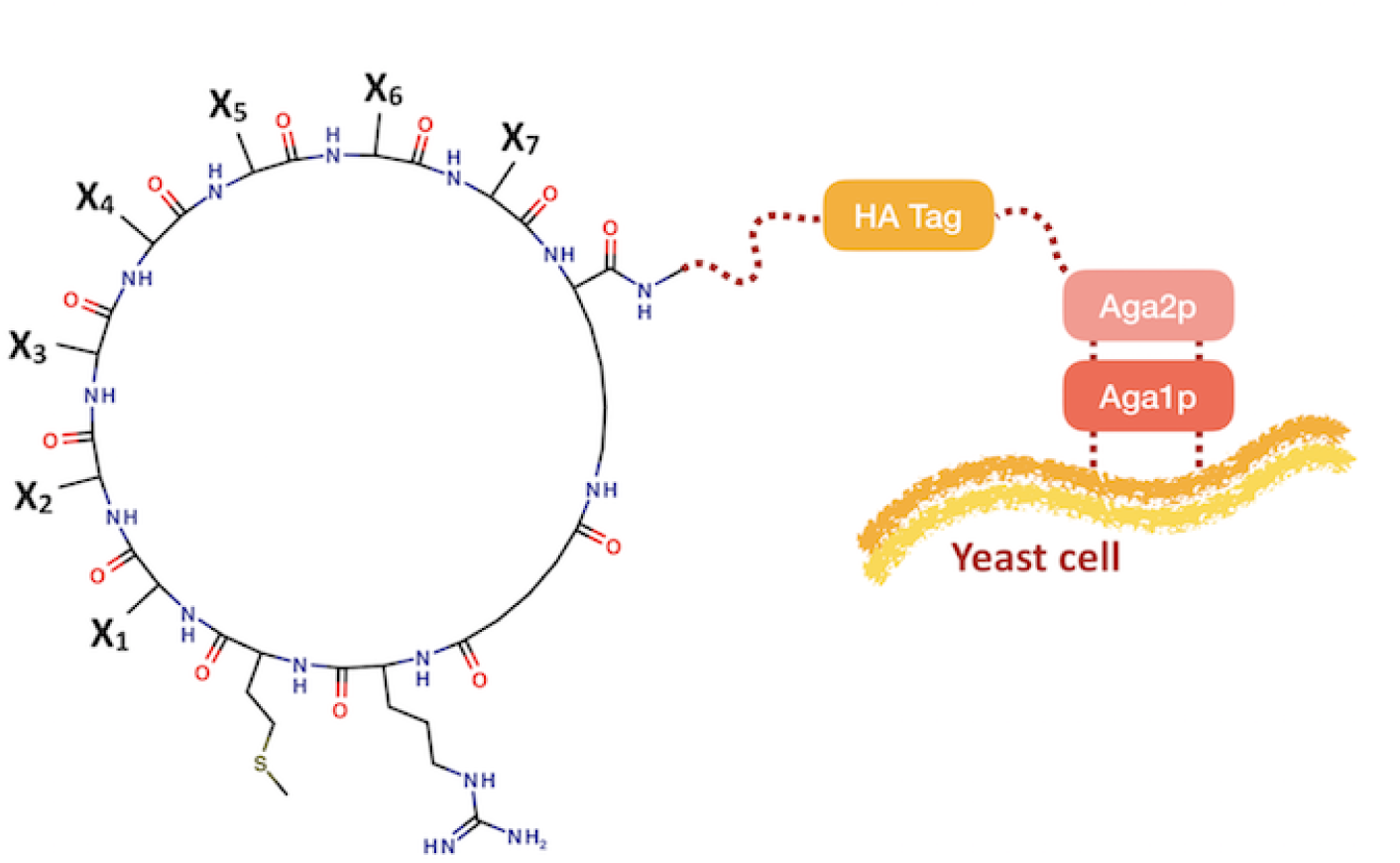
Yeast surface display of chemically crosslinked cyclic peptides. A linear peptide sequence is displayed as an N-terminal fusion to Aga2p, which is tethered to the yeast cell surface through a disulfide bond with Aga1p. The displayed peptides are crosslinked between the α-amine of arginine and the α-amine of a lysine located at the C-terminus of the peptide library’s variable region, using a DSG crosslinker.

To evaluate the use of yeast surface display for isolating cyclic peptide binders, we first screened a combinatorial library of chemically crosslinked cyclic peptides against a model target protein, lysozyme, using magnetic selection and FACS. We further evaluated if yeast displayed cyclic peptides can be used to efficiently assess binding specificity and to obtain quantitative estimates for equilibrium binding dissociation constants (K_D_), thereby bypassing the need to chemically synthesize isolated peptides for such evaluation. Finally, we investigated if our approach could be used to isolate and characterize cyclic peptides that bind human interleukin-17(IL-17).

## Results and Discussion

### Confirmation of peptide cyclization on the yeast surface

In our approach, the cyclization of a yeast-displayed linear peptide is mediated by a DSG cross-linker. While previously utilized to cyclize peptide fusions in solution^11,16^, to our knowledge, DSG has not been commonly used to cyclize peptide sequences expressed as cell surface fusions. Accordingly, we utilized a protein-binding assay to indirectly assess DSG-mediated cyclization of yeast-displayed linear peptide precursors. It is generally accepted that cyclic peptides exhibit higher target-binding affinity compared to their linear counterpart^17^. We hypothesized that if DSG appropriately cyclized linear displayed peptides then the target protein would bind more readily to DSG treated cells due to the higher affinity of cyclic peptides. Consequently, we compared the binding of human IgG to yeast displaying known IgG-binding cyclic peptide sequences in either their linear or DSG-cyclized form; the peptides cyclo-[*DSG*-MWFPHY-*K*], cyclo-[*DSG*-MHGFRG-*K*], and cyclo-[*DSG*-MWFRHY-*K*] had been selected in prior work to bind IgG^16^. Briefly, yeast cells displaying the linear IgG-binding peptides were incubated with the DSG cross-linker in two consecutive reactions to achieve peptide cyclization. To promote complete cyclization of all linear displayed peptides, DSG was incubated 550-fold over the total amount of peptides displayed on the yeast surface (~50,000 per cell^18^). Subsequently, DSG-treated and untreated cells were incubated with biotinylated IgG followed by immunofluorescent detection using Streptavidin R-phycoerythrin(SA-PE) (Figure 2). In comparison to the non-treated cells, binding of IgG to the DSG-treated cells was significantly higher, as evaluated by the mean fluorescence signal. This difference in binding is likely attributed to DSG appropriately crosslinking the linear displayed peptides resulting in higher affinity for IgG.

**Figure 2.**
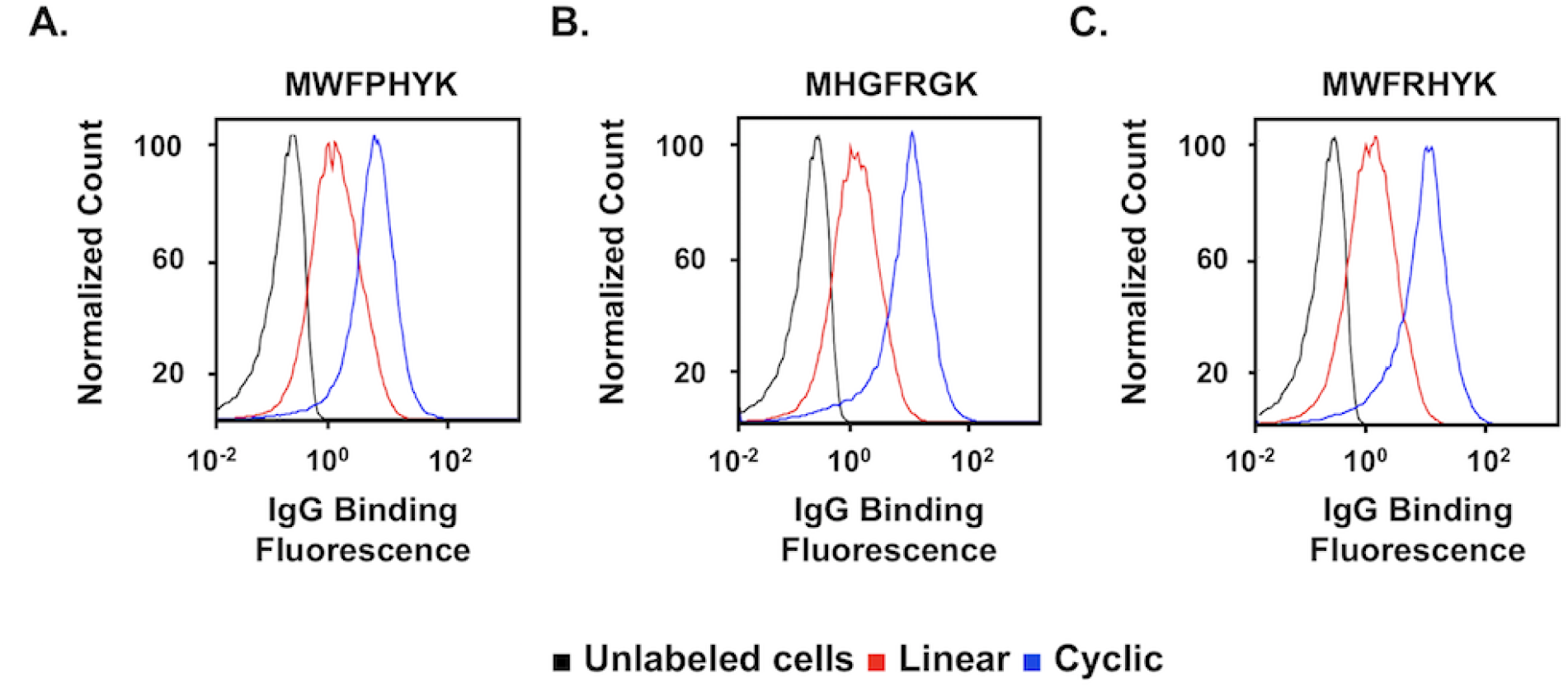
Binding of IgG to yeast cells displaying linear (red) and DSG crosslinked, cyclic (blue) peptide sequences compared to an unlabeled (no IgG) control (black). Three IgG specific cyclic peptide sequences were tested: **(A)** cyclo-[*DSG*-MWFPHY-*K*], **(B)** cyclo-[*DSG*-MHGFRG-*K*], and **(C)** cyclo-[*DSG*-MWFRHY-*K*]. Representative data from three independent experiments is shown. For each evaluated sequence, the mean fluorescence of IgG binding was significantly higher (p<0.05) for the DSG-treated cells (cyclic) compared to the non-treated cells (linear) using a two tailed, paired t-test.

### Isolation of lysozyme-binding cyclic peptides using yeast surface display

We generated a yeast-display library of linear peptide precursors with a diversity of ~ 5×10^7^ variants. The library variants were encoded as RMX_1_X_2_X_3_X_4_X_5_X_6_X_7_K, where X is a randomized amino acid residue. Subsequently,yeast displayed peptides were cyclized using DSG, as described earlier. We initially screened the library against a model target protein, lysozyme, using one round of magnetic selection with lysozyme-coated magnetic beads, followed by two rounds of FACS. Interestingly, we observed that simultaneous labeling of yeast cells with both lysozyme and an anti-HA antibody resulted in a decrease of lysozyme binding, relative to the case when the anti-HA antibody was not present (**Figure S1**). The binding of lysozyme to yeast-displayed cyclic peptides is likely affected by steric hindrance if the anti-HA antibody is bound. Therefore, unlike typical protocols used for theFACS selection of yeast display libraries, the cyclic peptide library was solely labeled with the target protein.

After FACS, plasmid DNA was isolated from 13 different colonies and sequenced (Table 1). Sequence homology, especially the presence of arginine (R) at position X_7_, and the enrichment in cationic (R and K), aromatic (W, F, and Y), and hydrogen-bonding (Q and S) residues is apparent in the selected sequences. The cationic character of the sequences is surprising given that lysozyme is a basic protein (pI~11^19^).

**Table 1.** Lysozyme binding peptides isolated from a yeast displayed library of cyclic heptapeptides. Mutagenized positions are denoted with an X. Numbers in parentheses represent the number of times a sequence occurred.

|  | M | X<sub>1</sub> | X<sub>2</sub> | X<sub>3</sub> | X<sub>4</sub> | X<sub>5</sub> | X<sub>6</sub> | X<sub>7</sub> | K |
| --- | --- | --- | --- | --- | --- | --- | --- | --- | --- |
| **LYS.1 (4)** |  | W | W | R | W | V | Y | R |  |
| **LYS.2 (3)** |  | I | C | W | F | C | N | K |  |
| **LYS.3 (2)** |  | R | Q | Y | R | S | R | R |  |
| **LYS.4** |  | W | W | R | L | V | Y | R |  |
| **LYS.5** |  | R | R | Q | G | L | R | R |  |
| **LYS.6** |  | R | R | R | R | P | V | R |  |
| **LYS.7** |  | K | R | F | K | T | R | R |  |

### Binding affinity analysis of lysozyme cyclic peptides

Lysozyme binding isotherms were generated for yeast cells displaying cyclo-[*DSG*-RMWWRWVYR*-K*], cyclo-[*DSG*-RMICWFLCN*-K*], cyclo-[*DSG*-RMRQYRSRR*-K*], cyclo-[*DSG*-RMRRQGLRR*-K*], and cyclo-[*DSG*-RMWWRLVYR*-K*]. Yeast cells displaying these cyclic peptides were incubated with varying concentrations of biotinylated lysozyme (20 nM - 8 μM). The fraction of cell surface fusions bound by biotinylatedlysozyme was quantified using immunofluorescent detection of SA-PE. The resulting data was fit using a monovalent binding isotherm (Figure 3A-E), to estimate the apparent binding affinity (K_D,Yeast_) for lysozyme (Figure 3F).

**Figure 3.**
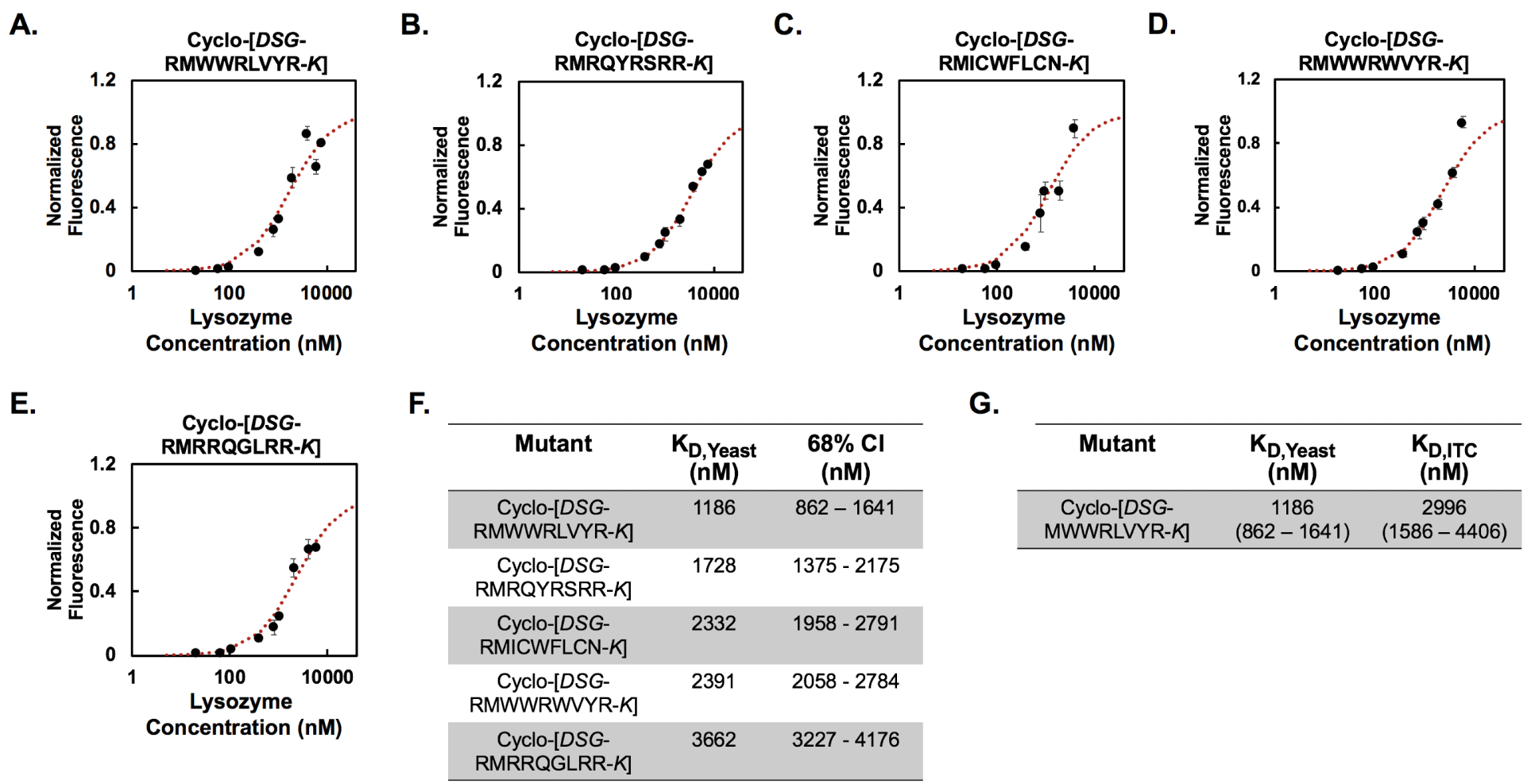
Estimation of apparent K_D_ describing the affinity of lysozyme for the yeast displayed cyclic peptides, **(A)** cyclo-[*DSG*-RMWWRLVYR-*K*], **(B)** cyclo-[*DSG*-RMRQYRSRR-*K*], **(C)** cyclo-[*DSG*-RMICWFLCN-*K*], **(D)** cyclo-[*DSG*-RMWWRWVYR-*K*], and **(E)** cyclo-[*DSG*-RMRRQGLRR-*K*]. Error bars represent the standard error of the mean for three independent experiments. **(F)** A global fit was used to estimate K_D,Yeast_ describing the affinity between the yeastdisplayed cyclic peptides and lysozyme as well as the corresponding 68% confidence interval (CI). **(G)** The binding affinity (K_D,ITC_) of soluble cyclo-[*DSG*-MWWRLVYR-*K*] for lysozyme was also estimated using isothermal calorimetry and compared to K_D,Yeast_. In parentheses, a 68% confidence interval is provided for K_D,Yeast_ while a range representing the standard deviation of the mean for four independent experiments is described for K_D,ITC_.

Cyclo-[*DSG-*RMWWRLVYR*-K*] exhibited the highest affinity for lysozyme. Accordingly, this cyclic peptide was chemically synthesized and characterized via isothermal calorimetry (ITC). The binding affinity of cyclo-[*DSG-*MWWRLVYR*-K*] for lysozyme as measured by ITC (2996 nM +/-1410 nM) was comparable to that obtained via yeast surface titration (862 – 1641 nM, 68% confidence interval) (Figure 3G). This suggests yeast surface titrations can be used to estimate the binding affinity of cyclic peptides, similar to proteins^18^. Quantitative assessment of binding affinity using yeast displayed cyclic peptides is particularly attractive as this method can be high throughput and circumvents the need to chemically synthesize peptides. It is important to note that the chemically synthesized peptides lack the N-terminal arginine residue that is present in the yeast displayed peptides. Removing arginine from the sequence was not expected to significantly alter the binding affinity of the peptide, given that a N-terminal arginine is present in all library peptides. Further, removing arginine, which displays a bulky side chain residue, considerably increases the yield of cyclization during synthesis.

The low-micromolar affinity of the lysozyme binding peptides was anticipated based on the affinity of previously identified cyclic peptides that are similar in size and amino acid composition^20^. Other groups, using display-based technologies, have identified cyclic peptides with low-nanomolar affinities^21^; these cyclic peptides, however, feature a larger ring structure (up to 14 randomized amino acid positions), and therefore benefit froma higher enthalpic contribution to binding free energy. In other instances, the peptide macrocycle is constrained via multifunctional crosslinkers to increase both the enthalpic and entropic contributions of binding free energy^22^.

### Specificity of cyclic peptides for lysozyme and effect of peptide cyclization on binding affinity

To assess binding specificity for lysozyme, we incubated yeast cells displaying the isolated cyclic peptides with varying concentrations (0.1 - 2 μM) of lysozyme or bovine serum albumin (BSA). Lysozyme binding was significantly higher than binding to BSA for all cyclic peptides (Figure 4A-E), demonstrating specificity for lysozyme. BSA was adopted as a model protein to test binding selectivity owing to its anionic character (pI ~5.4)^23^and because it was included in the buffers used during library screening. It is therefore notable that the selected peptides, despite their strong cationic and amphiphilic character, showed no or minimal BSA binding, supporting the claim that the proposed library screening method affords selective protein binders.

**Figure 4.**
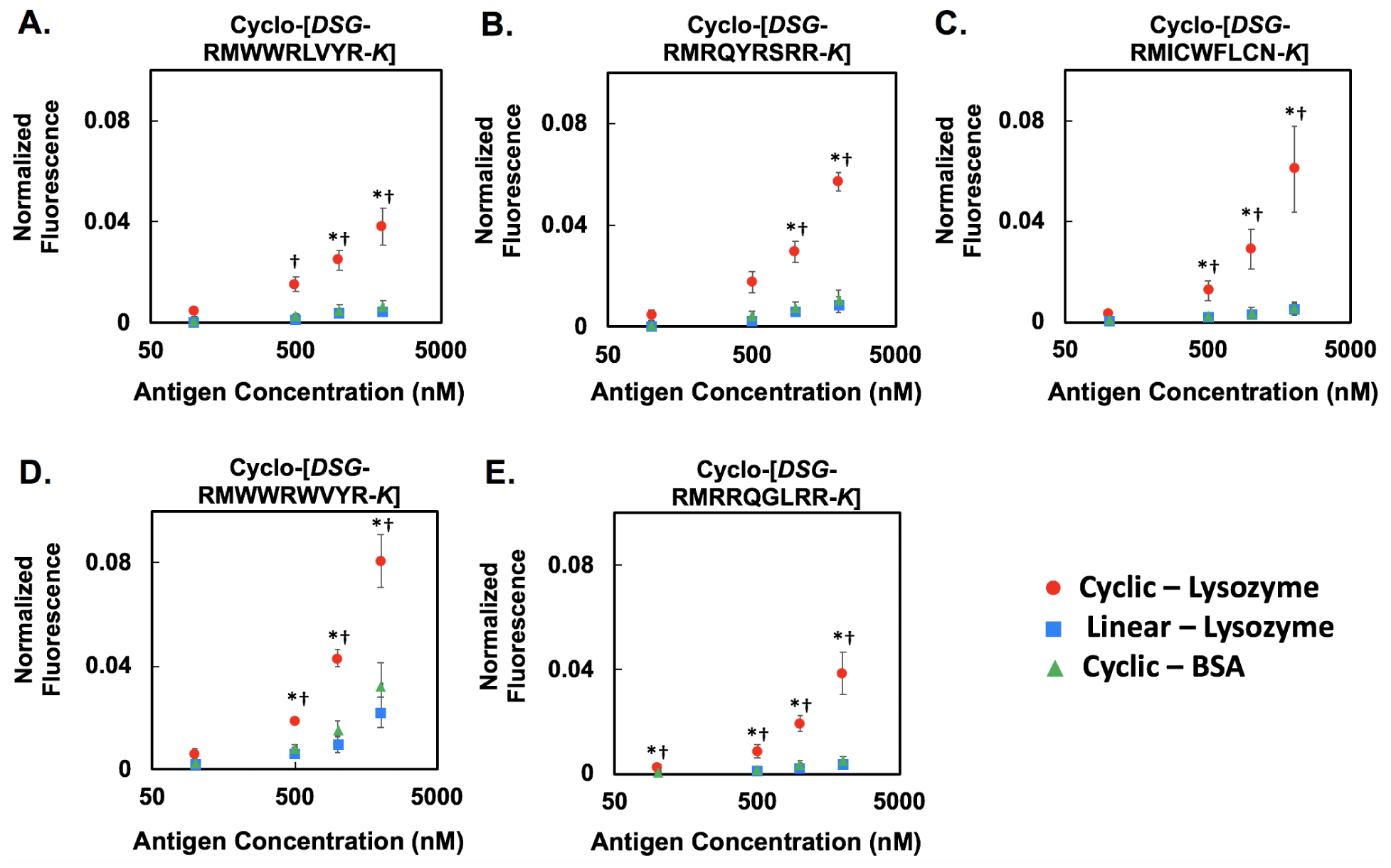
Specificity analysis of lysozyme binding cyclic peptides. The binding of lysozyme to yeast cells displaying cyclic peptides (red), **(A)** cyclo-[*DSG*-RMWWRLVYR-*K*], **(B)** cyclo-[*DSG*-RMRQYRSRR-*K*], **(C)** cyclo-[*DSG*-RMICWFLCN-*K*],**(D)** cyclo-[*DSG*-RMWWRWVYR-*K*], and **(E)** cyclo-[*DSG*-RMRRQGLRR-*K*], was compared to binding of lysozyme to yeast cells displaying their linear counterparts (blue). Additionally, the binding of lysozyme (red) was compared to the binding of BSA (green) for a given yeast displayed cyclic peptide. Error bars represent the standard error of the mean for three independent experiments. For a given lysozyme concentration, * represents statistically significant lysozyme binding (p< 0.05) for a yeast displayed cyclic peptide in comparison to its linear counterpart. At a specific antigen concentration, † represents statistically significant lysozyme binding (p< 0.05) in comparison to the binding of BSA for a given yeast displayed cyclic peptide. A two-tailed, paired t-test was performed to evaluate statistical significance.

We also evaluated the effect of peptide cyclization on lysozyme binding activity by comparing the extent of lysozyme binding (0.1 - 2 μM) to yeast cells expressing either the linear or cyclized form of each peptide. Lysozyme binding was significantly higher for the yeast cells displaying the cyclic form of each peptide (Figure 4A-E). All linear peptides, with the exception of RMWWRWVYRK (Figure 4D), displayed backgroundlevels of lysozyme binding for concentrations as high as 2 μM. This indicates that DSG-mediated cyclization critically confers binding activity to the selected sequences, and that biorecognition activity of cyclized peptides cannot be inferred based on their linear counterpart.

### Discovery of IL-17-binding cyclic peptides

To demonstrate broader applicability, we screened the cyclic heptapeptide library against Interleukin-17A (IL-17), a proinflammatory cytokine. The interaction of IL-17 with its receptor, IL-17RA, results in the stimulation of inflammatory signaling. Several autoimmune diseases, like plaque psoriasis and psoriatic arthritis, are characterized by excessive levels of IL-17^24^. Cyclic peptides specific to IL-17 can potentially be used as labeling agents^25^or inhibitors of the IL-17:IL-17RA interaction^26^.

Cyclic peptide binders for IL-17 were isolated from the yeast display heptapeptide library using magnetic selection and FACS. The magnetic selection was performed using magnetized yeast cells displaying IL-17 in lieu of target functionalized magnetic beads^27^. The use of a yeast-displayed target circumvents the need for purified target protein. The population resulting from FACS was subjected to Illumina Next Generation Sequencing (NGS). The isolated sequences (Table 2) are enriched in hydrophobic and basic residues, especially tryptophan (W) and arginine (R). The latter was not anticipated given the cationic character of IL-17 (sequence-based pI of 8.82).

**Table 2.** Sequences of IL-17 binders isolated from a yeast displayed library of cyclic heptapeptides. Mutagenized positions are denoted with an X. Numbers in parentheses represent the number of times a sequence appeared during Illumina Next Generation Sequencing.

|  | M | X<sub>1</sub> | X<sub>2</sub> | X<sub>3</sub> | X<sub>4</sub> | X<sub>5</sub> | X<sub>6</sub> | X<sub>7</sub> | K |
| --- | --- | --- | --- | --- | --- | --- | --- | --- | --- |
| **IL17.1 (4191)** |  | R | W | L | R | G | R | R |  |
| **IL17.2 (3446)** |  | L | F | H | W | P | F | P |  |
| **IL17.3 (2914)** |  | S | Y | F | W | R | W | L |  |
| **IL17.4 (2906)** |  | A | W | E | W | L | W | K |  |
| **IL17.5 (2742)** |  | L | F | F | D | W | L | L |  |
| **IL17.6 (2324)** |  | R | V | P | W | L | W | L |  |
| **IL17.7 (2099)** |  | W | R | N | F | A | W | R |  |
| **IL17.8 (2030)** |  | F | P | W | V | Q | W | R |  |
| **IL17.9 (1895)** |  | W | L | N | D | L | W | R |  |
| **IL17.10 (1891)** |  | F | L | W | H | F | R | T |  |

### In silico analysis of IL-17: cyclic peptide binding

Seventeen selected peptides **(Table S1)** were generated using atomistic molecular dynamic (MD) simulations and docked against IL-17 (PDB ID: 4HSA)^28^. It is important to note that the simulated peptide sequences lacked a N-terminal arginine residue. The IL-17-binding linear peptide IHVTIPADLWDWINK^26^ and cyclo-[*DSG-*MGSGGGGSG-*K*] were are also docked against IL-17 as positive and negative controls, respectively. Representative IL-17: peptide complexes selected from the top docking clusters were refined via MD simulations to estimate the free energy of binding (ΔG_b_) and the corresponding values of affinity (K_D,*in silico*_) (**Table S2**). Twelve of the tested cyclic peptides were predicted to form stable complexes with IL-17, featuring K_D,*in silico*_ ~ 23 nM – 1 μM (Figure 5A, **Figure S2**). Each of the peptides docked at the interface between IL-17 and IL-17R, suggesting these peptides may have the potential to modulate this interaction. As anticipated, the positive control peptide was predicted to bind IL-17 with a high affinity, K_D,*in silico*_ ~ 49 nM, which is reasonably similar to its experimentally predicted K_D_ (~2.6 nM)^26^. Similarly, the negative control cyclic peptide returned a K_D,*in silico*_ of 80 µM. Select peptide sequences were chosen for further *in vitro* evaluation, as they were predicted to bind IL-17 with affinities comparable to the positive *in silico* control.

**Figure 5.**
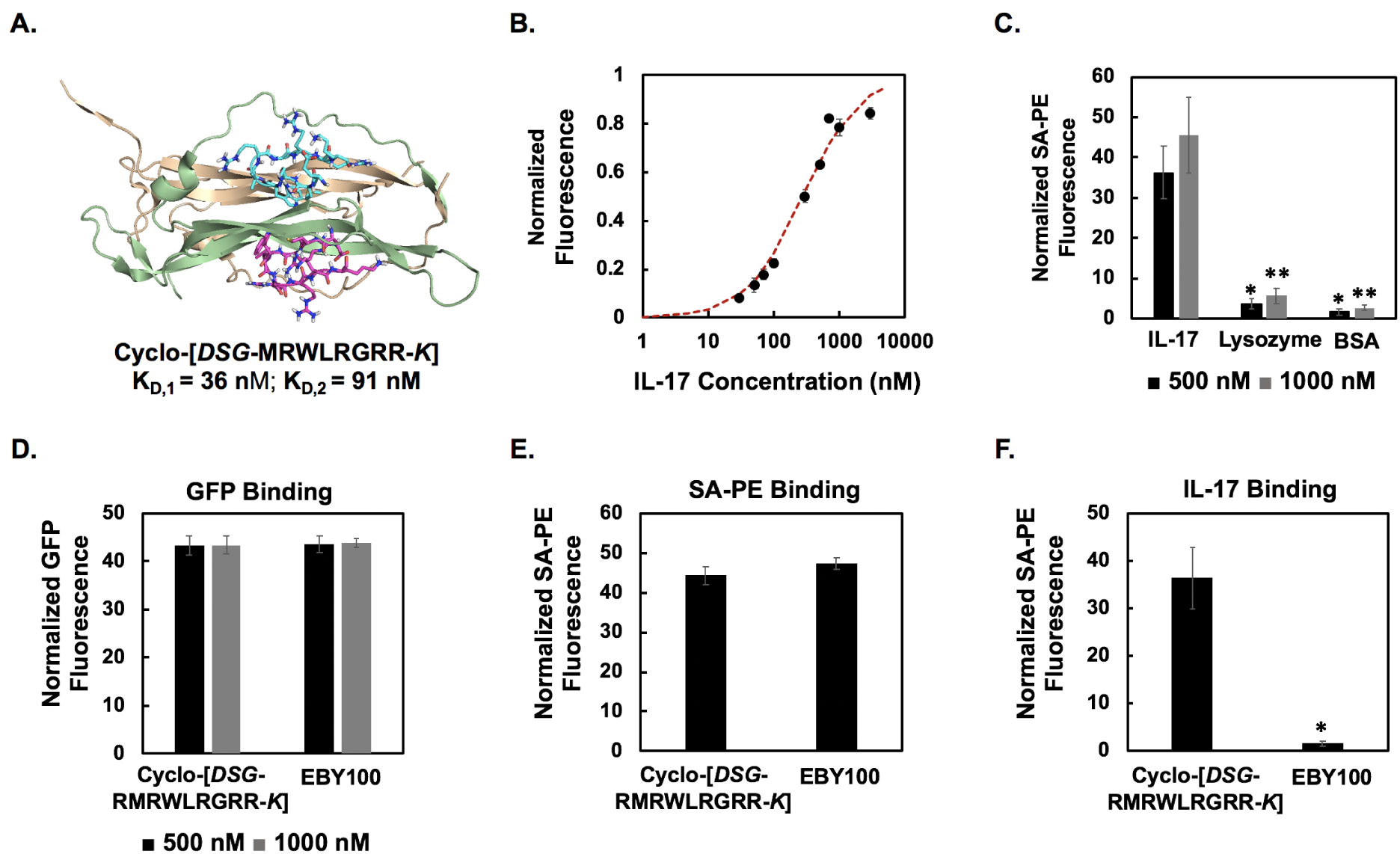
Affinity and specificity analysis of cyclo-[*DSG*-RMRWLRGRR-*K*] for IL-17. **(A)** *In silico* IL-17: cyclo-[*DSG*-MRWLRGRR-*K*] complex obtained by molecular docking and refined by molecular dynamic simulations. The monomers of IL-17 are denoted in green and brown while cyclo-[*DSG*-MRWLRGRR-*K*] is modeled in cyan or magenta. **(B)** Estimation of apparent *K*_*D*_ describing the affinity between IL-17 and yeast displayed cyclo-[*DSG*-RMRWLRGRR-*K*]. A global fit estimated the*K*_*D*_ as 279 nM (243 – 320 nM, 68% confidence interval). **(C)** Normalized mean fluorescence of IL-17, lysozyme, and BSA binding to yeast displayed cyclo-[*DSG*-RMRWLRGRR-*K*] at 500 and 1000 nM. Binding of these biotinylated proteins was detected using SA-PE. * denotes statistical significance in comparison to IL-17 binding at 500 nM for p<0.05 while ** denotes statistical significance in comparison to IL-17 binding at 1000 nM for p<0.05. **(D)** Normalized mean fluorescence of GFP binding to yeast displayed cyclo-[*DSG*-RMRWLRGRR-*K*] and DSG treated, non-displaying EBY100 yeast at 500 and 1000 nM. **(E)** Normalized mean fluorescence of SA-PE binding to yeast displayed cyclo-[*DSG*-RMRWL-RGRR-*K*] and DSG treated, non-displaying EBY100 yeast at 500 nM. **(F)** Normalized mean fluorescence of IL-17 binding to yeast displayed cyclo-[*DSG*-RMRWLRGRR-*K*] and DSG treated, non-displaying EBY100. Binding of biotinylated IL-17 was detected using SA-PE. * represents statistical significance (p< 0.05) in comparison to the binding of IL-17 to yeast cells displaying cyclo-[*DSG*-RMRWLRGRR-*K*]. Error bars represent the standard error of the mean for three individual replicates throughout. Statistical significance was evaluated using a two-tailed, paired t-test throughout.

### Binding affinity and selectivity of selected cyclic peptides for IL-17

The binding of IL-17 to yeast cells displaying peptides cyclo-[*DSG-*RMRWLRGRR-*K*], cyclo-[*DSG-*RMI-GQWWRR-*K*], cyclo-[*DSG-*RMNRLKFWF-*K*], cyclo-[*DSG-*RMSFFDIWR-*K*], cyclo-[*DSG-*RMYR-FHRHG-*K*], and cyclo-[*DSG-*RMFGLLHRG-*K*] was evaluated at a fixed concentration (500 nM). No significant difference in binding was observed between the various mutants (**Figure S3**). Hence, cyclo-[*DSG-*RMRWLRGRR-*K*] was chosen for subsequent evaluation as this sequence appeared most frequently in the NGS results. The affinity of cyclo-[*DSG-*RMRWLRGRR-*K*] for IL-17 was evaluated using the yeast surface titration method (Figure 5B). The K_D_ was estimated as 279 nM (243 – 320 nM, 68% confidence interval), which compares well with the predicted K_*D,in silico*_ of 36 - 91 nM; the small difference between the two values canbe attributed to binding non-ideality of the peptides when displayed on the yeast surface.

The binding selectivity of cyclo-[*DSG*-RMRWLRGRR-*K*] was evaluated by comparing the binding of IL-17 and multiple putative non-specific proteins to yeast cells displaying this peptide. Our results indicate that IL-17 bound to yeast-displayed cyclo-[*DSG*-RMRWLRGRR-*K*] at a significantly higher level (p<0.05) than lysozyme (pI~11^19^) and BSA (pI ~5^23^) (Figure 5C). While poor binding of lysozyme can be imputed to the electrostatic repulsion from the cationic cyclic peptide, it should be noted that IL-17 readily bound despite its mild cationic character (sequence-based pI of 8.82), indicating true binding affinity. The binding selectivity of the peptide against BSA, while engineered through the inclusion of BSA in the buffers utilized for library screening, is nonetheless notable, given the potential of BSA binding solely due to Coulomb interaction. Finally, green fluorescence protein (GFP) (pI~5.5^29^) and phycoerythrin (pI~4.2^30^) conjugatedstreptavidin (pI~5-6^31^) were found to bind similarly to cells expressing cyclo-[*DSG*-RMRWLRGRR-*K*] and non-displaying EBY100 (Figure 5D-E), suggesting that cyclo-[*DSG*-RMRWLRGRR-*K*] has no affinity for these proteins.

Lastly, we confirmed that IL-17 exhibits no interactions with non-specific proteins displayed on the surface of DSG-treated yeast. The large excess of DSG employed during the cyclization of linear displayed peptides is likely to promiscuously modify other yeast surface proteins. The possibility of IL-17 interacting with DSG-modified surface proteins cannot be excluded *a priori*. To evaluate this, we compared the binding of IL-17 to DSG-treated null EBY100 cells *vs.* peptide-displaying yeast cells (Figure 5F). The lack of binding to the null cells corroborates the claim that cyclo-[*DSG*-RMRWLRGRR-*K*] specifically interacts with IL-17 and provides confidence that off-target DSG modifications do not influence the identification of peptide binders during library screening.

### Evaluation of cyclo-[DSG-MRWLRGRR-K] as a modulator of the IL-17: receptor interaction

While the specificity of cyclo-[*DSG*-RMRWLRGRR-*K*] for IL-17 supports the use of this peptide as an affinity ligand, we explored the use of cyclo-[*DSG*-MRWLRGRR-*K*] to modulate the interaction between IL-17 and its receptor, as suggested by *in silico* modeling (Figure 6A). Peptides and other small molecules have been used to modulate protein-protein interactions both as inhibitors and stabilizers^32^. Specific to the IL-17: receptor interface, linear peptide IHVTIPADLWDWINK was previously developed as inhibitor of this interaction for the potential use as an anti-inflammatory^26^. No specific stabilizers of the IL-17: IL-17RA interaction have been developed, although a stabilizing molecule that increases the affinity between IL-17 and its receptor could have implications for mesenchymal stem cell proliferation and differentiation, which have been shown to be dependent on this interaction^33^. Accordingly, we performed an *in vitro* evaluation to understand if cyclo-[*DSG-*MRWLRGRR-*K*] modulates the interaction between IL-17 and IL-17RA.

**Figure 6.**
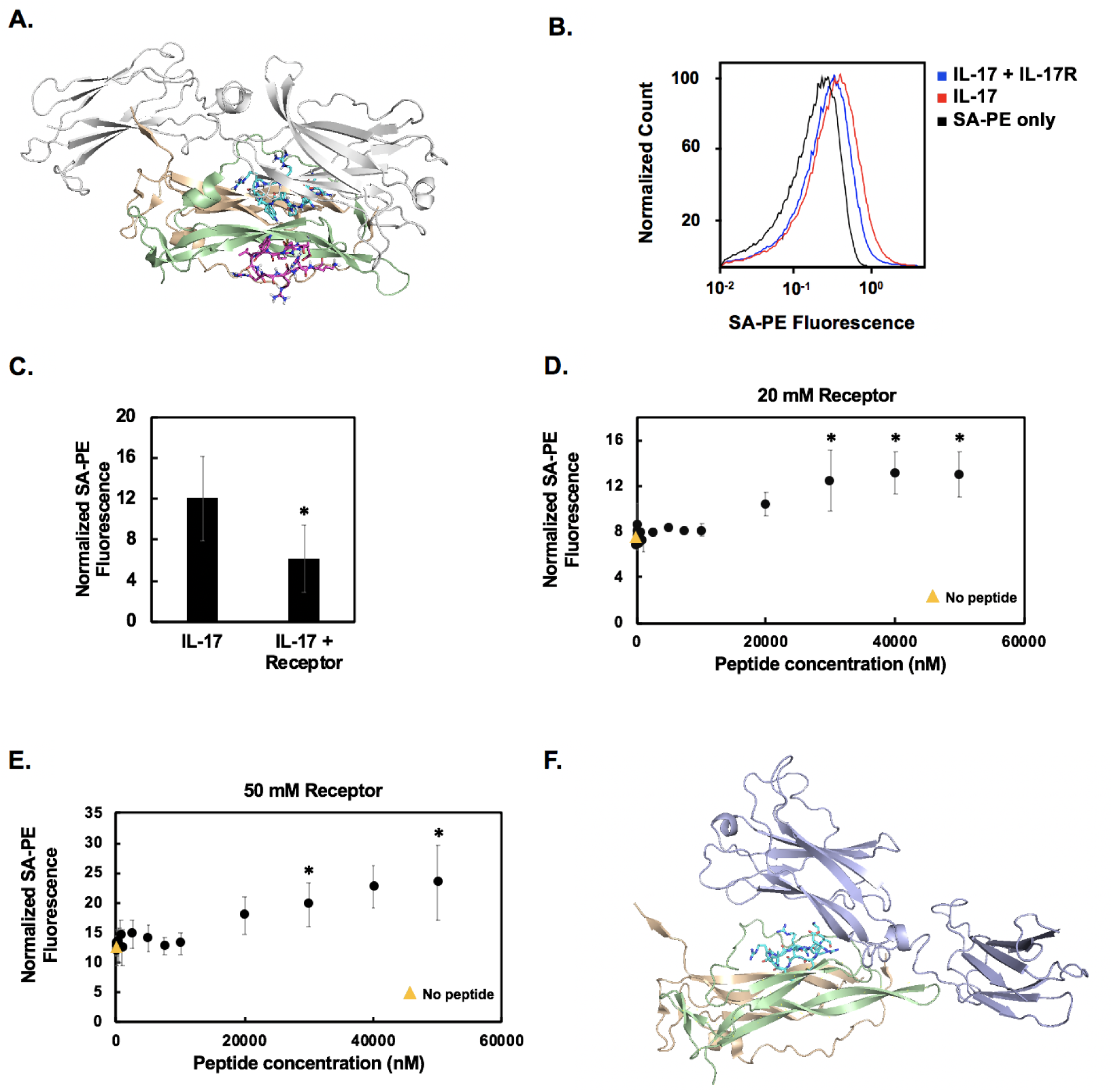
Evaluation of cyclo-[*DSG*-MRWLRGRR-*K*] as a potential modulator of IL-17’s interaction with its receptor (IL-17RA). **(A)** *In silico* modeling of cyclo-[*DSG*-MRWLRGRR-*K*] (cyan or magenta) interacting with IL-17 (monomers denoted in green and brown). IL-17RA was not present during the simulation. Rather, the interaction of IL-17RA with IL-17 was superimposed after modeling and is visualized in grey. This visualization suggests cyclo-[*DSG*-MRWLRGRR-*K*] interacts with IL-17 in the same region as IL-17RA. **(B)** Yeast cells displaying cyclo-[*DSG*-RMRWLRGRR-*K*] were incubated with 100 nM biotinylated IL-17 in the presence (blue) or absence (red) of 500 nM IL-17RA followed by SA-PE detection. Yeast cells displaying cyclo-[*DSG*-RMRWLRGRR-*K*] were also incubated with only SA-PE (black). A representative histogram of three repeats is shown. **(C)** The normalized mean fluorescence of IL-17 binding to the yeast displayed cyclo-[*DSG*-RMRWLRGRR-*K*] was lower when IL-17RA was present (* denotes p<0.001 for a two tailed, two sample unequal variance t-test). This suggests that cyclo-[*DSG*-RMRWLRGRR-*K*] binds in a similar region to IL-17 as IL-17RA. The effect of peptide concentration on the modulation of IL-17’s interaction with its receptor is detailed in **(D)** and **(E)**. Yeast cells displaying IL-17 were incubated with soluble cyclo-[*DSG*-MRWLRGRR-*K*] and IL-17RA. The binding of IL-17RA was detected using biotinylated protein A and SA-PE. The concentration of peptide was varied from 0 nM to 50 µM while the concentration of receptor was held at either 20 nM **(D)** or 50 nM **(E)**. The binding of IL-17RA when no peptide was present is denoted with a yellow triangle for comparison. * represents a statistical difference (p<0.10 for a two tailed, paired t-test) of IL-17RA binding in comparison to when no peptide is present. **(F)** *In silico* IL-17: cyclo-[*DSG*-MRWLRGRR-*K*]: IL-17RA complex. All three molecules were present during the simulation. IL-17’s individual monomers are denoted in green and brown while IL-17RA is simulated in purple. cyclo-[*DSG*-MRWLRGRR-*K*] is modeled in cyan. Error bars represent the standard error of the mean for three individual replicates throughout.

To this end, a two-step evaluation approach was implemented. First, yeast cells displaying cyclo-[*DSG-*RMRWLRGRR-*K*] were incubated with soluble, biotinylated IL-17 (100 nM) in the presence or absence of IL-17RA (500 nM). The binding of biotinylated IL-17 to the cells was measured by flow cytometry using SA-PE for detection (Figure 6B). The fluorescence signal corresponding to the binding of IL-17 was slightly lower(p<0.001) in presence of IL-17RA (Figure 6C), corroborating the *in silico* observation that cyclo-[*DSG-*RMRWLRGRR-*K*] and IL-17RA may bind to IL-17 in a similar region (Figure 6A). Importantly, we have shown that cyclo-[*DSG-*RMRWLRGRR-*K*] does not interact with IL-17RA (**Figure S4**).

For the second approach, soluble cyclo-[*DSG-*MRWLRGRR-*K*] and soluble IL-17RA were incubated with yeast-displayed IL-17 to evaluate the effects of peptide concentration on IL-17RA binding to IL-17. The binding of IL-17RA to the yeast-displayed IL-17 was detected using biotinylated protein A and SA-PE; note that the IL-17RA used in this work is a Fc fusion. The concentration of IL-17RA was maintained at 20 or 50 nM, while the peptide concentration varied from 0 - 50 µM. The binding of IL-17RA to yeast displayed IL-17 was not significantly altered for peptide concentrations less than 30 µM, indicating no inhibition of the IL-17:IL-17RA interaction (Figure 6D-E). While cyclo-[*DSG-*MRWL-RGRR-*K*] likely shares an IL-17 binding site with IL-17RA, as suggested by the first *in vitro* modulation experiment, no inhibition was observed in this second experiment due to the superior IL-17-binding strength of IL-17RA (K_D_~2.8 nM)^34^ compared to that of the peptide (K_D_~279 nM) as well as the high avidity of binding between the bivalent IL-17RA Fc fusion and yeast displayed IL-17. Interestingly, the binding of IL-17RA to IL-17, increased slightly (p<0.1) for high concentrations (≥30 µM) of the cyclic peptide in comparison to when no peptide was present. We explored this phenomenon using MD simulations (Figure 6F), which returned a slightly lower K_D,*in silico*_ (higher affinity) for IL-17:cyclo-[*DSG-*MRWL-RGRR-*K*]:IL-17RA (14 nM) than for IL-17:IL-17RA (52 nM). These results suggest that cyclo-[*DSG-* MRWLRGRR-*K*] may act as a potential stabilizer of the IL-17:IL-17RA interaction. It is possible that the conformation of IL-17 changes as a result of peptide binding, thereby increasing IL-17’s affinity for IL-17RA. However, further *in vitro* analysis is needed to investigate this effect.

Collectively, our results demonstrate the successful development of a method for the efficient identification and characterization of cyclic peptide ligands with selective protein-binding activity. Of note is the integration of FACS, which accelerates the library screening process. The use of yeast displayed cyclic peptides allows on-the-fly specificity and affinity evaluation of potential peptide ligands. We envision this method will enable the identification of cyclic peptides to be used as affinity ligands for protein purification^16^ and biosensor^35^ applications that require moderate affinity biorecognition moieties as well cyclic peptides that inhibit moderate affinity protein-protein interactions.

## Materials and Methods

### Cyclization of linear peptides on the yeast surface

1×10^8^ yeast cells expressing a linear peptide sequence were washed 3X with crosslinking buffer (0.2 M NaCl, 1 mM EDTA). The cells were resuspended in 2 mL of crosslinking buffer and 10 µL of DSG cross-linker (Thermo Fisher Scientific) dissolved in N,N’-dimethylformamide (DMF) at 1.5 mg/mL. The cross-linking reaction was performed for 2 hours at 4°C. After, the cells were washed 3X with crosslinking buffer, and the crosslinking reaction was repeated. Finally, the cells were washed three times with 50 mM Tris, 300 mM NaCl pH 7.4 and stored in this buffer overnight prior to subsequent use.

### Binding analysis to confirm cyclization of yeast displayed linear peptides

The interaction of IgG with yeast cells displaying the linear or cyclized forms of each IgG binding peptide was characterized via flow cytometry. 5×10^6^ cells were labeled with 450 nM biotinylated IgG (Immunoreagents) for15 minutes at room temperature. After, the cells were washed and labeled with a 1:250 dilution of SA-PE (Thermo Fisher Scientific) for 12 minutes on ice. Cells were analyzed using a Miltenyi Biotec MACsQuant VYB cytometer.

### Generation of a cyclic heptapeptide yeast display library

A yeast display library of linear peptides with seven randomized amino acid positions flanked between a methionine (N-terminus) and a lysine (C-terminus) was generated using the previously described lithium acetate yeast transformation method^36^. Detailed protocols are found in the supplemental methods.

### Screening a cyclic heptapeptide yeast display library

Library screening was conducted using magnetic sorting and FACS, as detailed in prior work^18^. The magnetic sort against lysozyme (Thermo Fisher Scientific) employed streptavidin beads functionalized with lysozyme,whereas magnetic sorting against IL-17 was performed using magnetic yeast displaying IL-17^27^. Additional details are provided in the supplemental methods.

### Binding affinity estimation using yeast surface titrations

The K_D_ describing the interaction between a target antigen and a yeast displayed peptide was estimated using yeast surface titrations^18^. Briefly, yeast cells displaying a particular peptide were incubated with varyingconcentrations of biotinylated antigen (*i.e.*, lysozyme or IL-17) followed by SA-PE detection. The K_D_ was estimated using a global fit analysis. Detailed protocols are found in the supplemental methods.

### Binding affinity estimation using isothermal calorimetry

The K_D_ of lysozyme and chemically synthesized cyclo-[*DSG-*MWWRLVYR*-K*] was predicted using a nano-ITC calorimeter (TA instruments). 500 μM of the peptide was injected into a 50 μM solution of lysozyme. Additional details are provided in the supplemental methods.

### Binding specificity characterization

The binding selectivity of the identified cyclic peptide mutants was analyzed by labeling yeast-displayed cyclic peptides with specific antigens followed by secondary immunofluorescent detection. Detailed protocols are found in the supplemental methods.

### In silico evaluation of the IL-17: cyclic peptide and IL-17:peptide:IL-17RA interactions

Equilibrated peptide crystal structures were docked *in silico* against IL-17 using HADDOCK^37,38^. Representative IL-17: peptide complexes selected from the top docking clusters were refined using MD simulations. The MM/GBSA method was used to estimate each complex’s free energy of binding and the corresponding values of affinity^39,40^. Similar methods were used to simulate the interaction of IL-17RA with the MD refined IL-17:peptide complexes. Additional simulation details are provided in the supplemental methods.

### In vitro evaluation of cyclo-[DSG-MRWLRGRR-K] as a modulator

Two approaches were implemented: 1) incubate yeast cells displaying cyclo-[*DSG-*RMRWLRGRR*-K*] with 100 nM of soluble biotinylated IL-17 (R&D systems) in presence or absence of 500 nM IL-17RA (R&D systems), or 2) incubate yeast cells displaying IL-17 with varying concentrations of soluble cyclo-[*DSG-*MRWLRGRR*-K*] (0 - 50 μM) in presence of IL-17RA (20 or 50 nM) overnight at 4ºC. The binding of IL-17 was detected by SA-PE labeling, whereas the binding of IL-17RA was detected via biotinylated-protein A (Thermo Fisher Scientific) and SA-PE labeling. Additional details are provided in the supplemental methods.

## Supporting information

Supplemental experimental methods, a comparison of FACS labeling methods, IL-17: peptide complexes predicted from molecular docking, a comparison of IL-17 binding to yeast displayed cyclicpeptides, an evaluation of IL-17RA binding to yeast-displayed cyclo-[*DSG*-RMRWLRGRR-*K*], a plasmid schematic for yeast display of linear peptides, a table of IL-17 binding peptides used in docking studies, a table of binding energies and calculated affinity dissociation constants for IL-17: peptide complexes predicted by molecular simulation, a table of gene blocks fragments, a table of oligonucleotide primers, and a table of oligonucleotide DNA fragments are provided.

## Acknowledgments

This work was funded by a grant from the National Science Foundation (CBET 1511227). KBB kindly acknowledges support from an NSF Graduate Research Fellowship and a National Institute of Health Molecular Biotechnology Training Fellowship (NIH T32 GM008776). We wish to thank the UNC High-Throughput Peptide Synthesis and Array facility for peptide synthesis.

---

For Table of Contents Only

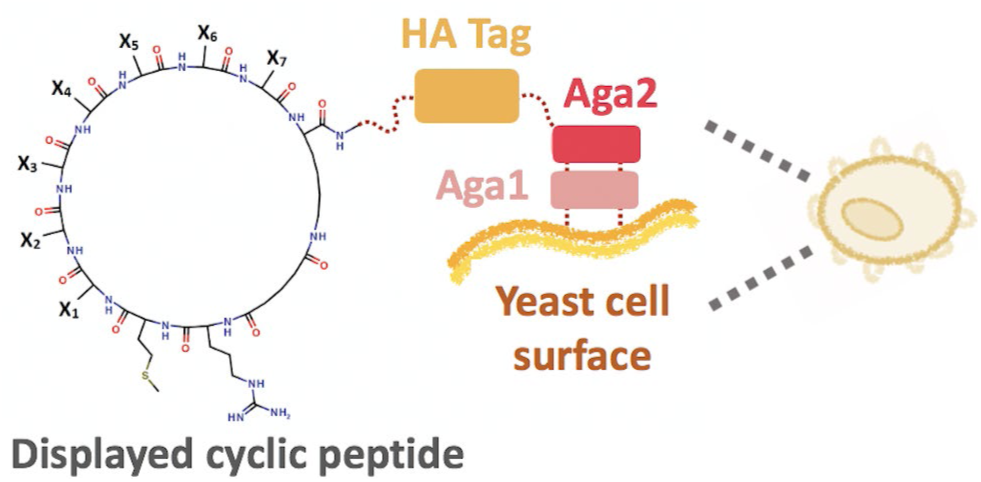

