## Supporting Information for "Isolation of Chemically Cyclized Peptide Binders Using Yeast Surface Display"

Kaitlyn Bacon[1], Abigail Blain[1], Matthew Burroughs[1], Nikki McArthur[1], Balaji M. Rao[1,2*], and Stefano Menegatti[1,2*]
[1]Department of Chemical and Biomolecular Engineering, North Carolina State University, NC, USA
[2]Biomanufacturing Training and Education Center (BTEC), North Carolina State University, NC, USA

**SUPPORTING INFORMATION**

**Contents:**

Supplementary Methods

5 Supplementary Figures

5 Supplementary Tables

**Supplementary Methods**

*Plasmids and yeast culture*

Cells were cultured using TRP-deficient SDCAA media for cells containing the pCTCON vector, and SGCAA medium was used to induce cell surface protein expression, as described previously[1]. Leu-deficient SDSCAA (-Leu) and SGSCAA (-Leu) media was similarly used for cells containing a pCT302-based plasmid[2]. Leu-deficient media is similar in composition to SDCAA and SGCAA media except it does not contain casamino acids, but rather synthetic drop out mix (1.62 g/L) lacking leucine (US Biological Life Sciences). Yeast cells were cultured in the appropriate growth media, SDCAA or SDSCAA, at 30°C with shaking at 250 RPM. To induce protein expression, yeast cells were transferred to SGCAA or SGSCAA medium at an $OD_{600}$ of 1 and cultured overnight at 20 °C with shaking at 250 rpm. EBY100 yeast without plasmid was grown in YPD medium (10.0 g/L yeast extract, 20.0 g/L peptone, and 20.0 g/L dextrose) at 30 °C with shaking at 250 rpm. Plasmid DNA was transformed into chemically competent *Saccharomyces cerevisiae* strain EBY100 using the Frozen-EZ Yeast Transformation Kit II (Zymo Research).

*Construction of peptide display plasmids*

pCTCON-Nterm was constructed by amplifying gene block 1 with Pf1 and Pr1 and introduced into pCTCON via the EcoRI and XhoI sites. This particular gene block displays a random linear peptide, between the NheI and BamHI restriction sites. To display specific linear peptide sequences on the yeast surface, pCTCON-Nterm was cut with NheI and BamHI, and DNA encoding a linear peptide sequence was introduced via homologous recombination. The linear peptide sequence DNA was purchased with the necessary homology arms as single stranded oligonucleotides and amplified using Pf2 and Pr2. These homology arms eliminate the NheI and BamHI sites present in pCTCON-Nterm **(Figure S5A)**. The linear sequence of three previously identified cyclic peptides[3], with affinity for IgG, were displayed on the yeast surface using oligo DNA blocks that encode the sequences MWFPHYK, MHGFRGK, and MWFRHYK, respectively.

All double stranded gene block fragments were purchased from Integrated DNA Technologies (IDT). All primer oligonucleotides and oligonucleotide DNA fragments were purchased from IDT or Eton Biosciences. Gene block fragments, oligonucleotide primers, and oligonucleotide DNA fragments can be found in **Tables S3-S5**. PCR reactions were performed at a 50 µL scale using Phusion polymerase (Thermo Fisher Scientific) according to the manufacturer's protocol. The amplified DNA was purified using the 9K Series Gel and PCR extraction kit from BioBasic. Restriction enzymes were bought from New England Biolabs. Plasmids and PCR products were digested in a 50 µL reaction volume with a 5X excess of each restriction enzyme for 4 hours at 37°C. Digested plasmid backbones were subsequently incubated with Antarctic Phosphatase purchased from New England Biolabs for 1 hour at 37°C. Digested plasmids and PCR products were purified using the BioBasic 9K Series Gel and PCR extraction kit. Ligations combining digested plasmid backbone and inserts were carried out overnight at 16°C using T4 DNA ligase (Promega) prior to transformation into electrocompetent Novablue *E. coli* cells. Transformations were carried out at 1.6 KV, 25 µF, and 200 Ω. The GeneJET plasmid miniprep kit (Thermo Fisher Scientific) was used to harvest DNA plasmids from overnight *E.coli* cultures.

When cloning using homologous recombination, 200 ng of cut plasmid and 250 ng of insert were transformed into chemically competent EBY100 cells prepared with the Frozen-EZ Yeast Transformation Kit II. The plasmid and insert were concentrated using ethanol precipitation such that the volume of DNA used in the transformation step was less than 5 µL in total. Briefly, a 1/10 volume of potassium acetate and 10 µL of linear acrylamide was added to the DNA to be concentrated followed by 2 volumes of icecold ethanol. The mixture was stored at -20 °C overnight. The next day, the DNA was centrifuged at 15,000 g for 10 minutes followed by removal of the supernatant. The DNA pellet was washed once with 70% ethanol followed by a wash with 100% ethanol and dried prior to resuspension in water. Recombined plasmid was extracted from the transformed yeast cells using the Zymoprep™ Yeast Plasmid Miniprep II kit following the manufacturer's recommendations. The recovered plasmids were transformed into electrocompetent Novablue *E. coli* cells, and the DNA was recovered from an overnight *E. coli* culture using the GeneJet plasmid miniprep kit.

*Generation of a cyclic heptapeptide yeast display library*

A yeast display combinatorial library of linear peptides was generated by amplifying gene block 2 with primers Pf3 and Pr3, that contain consensus sequences with pCT-NT-F2A-Sso7dhFc, in eight identical 50 µL reactions[4]. Gene block 2 encodes a linear peptide sequence that contains seven randomized amino acid positions flanked between a methionine and lysine. The peptide sequence is expressed as a N-terminal fusion to Aga2p **(Figure S5B)**. A HA tag follows the peptide sequence to confirm expression of the peptide on the yeast surface via immunofluorescence. The amplified product from gene block 2 was purified using phenol: chloroform extraction followed by DNA concentration using ethanol precipitation as previously described. Concurrently, 40 µg of a previously engineered plasmid, pCT-NT-F2A-Sso7dhFc, was linearized using EcoRI and XmaI[4]. The digested backbone was purified and concentrated using phenol: chloroform extraction and ethanol precipitation. The yeast library was generated using a previously described lithium acetate transformation protocol where the amplified product from gene block 2 and the digested backbone vector were transformed into EBY100 yeast via electroporation[5]. Three electroporations were performed using a Bio-Rad Gene Pulser system where 12 µg of PCR product and 4 µg of digested vector were combined with 400 µL of electrocompetent EBY100. The electroporation settings used were 2500 V, 25 µF, and 250 Ω. A vector only electroporation was performed as a control. The library diversity was determined by plating serial dilutions of the transformation reaction onto SDCAA plates and estimated as $5x10^7$.

*Screening a cyclic heptapeptide yeast display library against lysozyme*

The cyclic heptapeptide yeast display library was screened against lysozyme in one round of magnetic selection and two rounds of fluorescence activated cell sorting (FACS). To perform the magnetic selection, 25 µL of Biotin Binder Dynabeads (Thermo Fisher Scientific) were pre-coated with 10.8 µM biotinylated lysozyme overnight at 4°C in 100 µL of 0.1% BSA PBS (0.1% PBSA). Lysozyme (Thermo Fisher Scientific) was previously biotinylated by incubating a 3:1 molar excess of Ez-Link™Sulfo-NHS-LC-Biotin (Thermo Fisher Scientific) for two hours at 4°C. Excess reagent was removed by performing dialysis against 50 mM Tris-HCl, 300 mM NaCl pH 7.4. Protein concentrations were estimated using a BCA assay.

For selection, the library was induced and cyclized as described in the main methods section. The following day, a negative selection was performed where $5x10^8$ cyclized library cells were incubated with 25 µL of non-functionalized Biotin Binder Dynabeads at room temp for 1 hour. Cells that did not bind the non-functionalized beads were recovered and an additional negative selection was carried out using a new set of non-functionalized beads. Any unbound cells were subsequently removed and used in a positive selection against the pre-coated lysozyme beads for 1 hour at room temperature. Cells that did not bind the lysozyme functionalized magnetic beads were removed using a magnet. After, the lysozyme functionalized magnetic beads and any associated library cells were washed 3X with 0.1% PBSA, resuspended in SDCAA media, and incubated for 48 hours at 30°C to allow for expansion of the positively bound cell population. All selections took place in 4 mL of 0.1% PBSA. All beads were washed 3X with 0.1% PBSA prior to incubation with the library cells.

The pool of cells obtained from the magnetic selection were cyclized, and FACS was performed on this population using a MoFlo (Beckman Coulter) flow cytometer as previously described[1]. For FACS, $1x10^7$ cells were labeled with biotinylated lysozyme on ice for 1 hour. After, the cells were labeled with SA-PE (1:250 dilution) on ice in the dark for 12 minutes. All labelings took place in 100 µL of 0.1% PBSA. Washes with 0.1% PBSA were performed after each incubation. The cells were not double labeled for lysozyme binding and HA expression as the binding of the anti-HA antibody appeared to hinder the peptide's ability to interact with lysozyme. Two rounds of FACS were performed where the library cells were labeled with 1 µM of lysozyme for both rounds. After the second round of FACS, the isolated cells were plated on SDCAA plates (20 g/L dextrose, 5 g/L casamino acids, 6.7 g/L yeast nitrogen base, 182 g/L sorbitol, 5.4 g/L $Na_2HPO_4$, 8.6 g/L $NaH_2PO_4 \cdot H_2O$, 12 g/L Agar), and 13 clones were selected for sequencing. Plasmids were extracted from yeast cultures of the individual clones using a Zymoprep™ Yeast Plasmid Miniprep II kit. For each clone, the extracted plasmid DNA was transformed into electrocompetent Novablue cells and plated. An overnight *E.coli* culture was grown for each clone followed by DNA extraction and sequencing.

*Binding affinity estimation using yeast surface titrations*

The affinity ($K_D$) between the isolated cyclic peptides and their target (lysozyme or IL-17) was estimated using a yeast surface titration method. Briefly, yeast-displayed linear peptide sequences were cyclized using the technique described in the main methods section. Subsequently, the yeast cells displaying a cyclic peptide of interest were labeled with varying concentrations of the target protein, either biotinylated lysozyme (20 nM – 8 µM) or IL-17 (30 nM – 3 µM; R&D Systems), for 1 hour at room temperature. This was followed by secondary labeling with SA-PE (1:250 dilution) for 12 minutes on ice. The cells were then analyzed by flow cytometry as previously described[1]. All labeling took place in 50 µL of 0.1% PBSA. Washes with 0.1% PBSA were performed after each incubation. A SA-PE only labeling was also performed for yeast cells displaying each cyclic peptide to quantify the background mean fluorescence. The $K_D$ describing the binding affinity between a yeast displayed peptide and a target protein was estimated using the following relationship:

$$F = \frac{F_{Max}[L]_O}{K_D + [L]_O} \quad \text{(Eq. 1)}$$

Where $F$ is the background subtracted mean fluorescence, $[L]_0$ is the concentration of antigen used to label the cells, and $F_{Max}$ is the background subtracted fluorescence intensity when surface saturation is attained. The binding data was fit to Eq.1 using a global non-linear least squares regression across three repeats as previously described[1]. A single $K_D$ value and a unique $F_{Max}$ for each repeat were used as the fitted parameters. At high concentrations, the fluorescence signal decreased due to the Hook effect[6]. We ignored these points when applying the non-linear regression fit. Error bars represent the standard error of the mean for three independent experiments.

*Binding affinity estimation using isothermal calorimetry*

The affinity of cyclo-[*DSG*-MWWRLVYR-*K*] for lysozyme was estimated using a nano-ITC calorimeter (TA Instruments). A soluble form of this peptide was synthesized by the UNC High-Throughput Peptide Synthesis and Array Facility[3]. The soluble peptide lacks the N-terminal arginine residue that is present for the yeast displayed peptides. 500 µM of the peptide dissolved in water was injected into 50 µM of lysozyme dissolved in water. 20 injections of 2.5 µL were carried out 250 seconds apart at 25°C. The reference cell contained water. We performed this titration in quadruplicate. A blank titration was also performed where water was injected into 50 µM of lysozyme using the same conditions. The blank titration was only performed once. Binding isotherms were analyzed using software provided by TA instruments where the experimental titration values were normalized by the blank titration values.

*Specificity characterization of lysozyme-binding cyclic peptides*

The specificity of the lysozyme-binding cyclic peptides was evaluated using two different approaches. First, the binding of lysozyme to yeast cells displaying the cyclic form of a peptide was compared to the binding of lysozyme to yeast cells displaying the linear counterpart. Briefly, $2x10^6$ cells displaying either the linear or cyclic form of each peptide were incubated with 100, 500, 1000, or 2000 nM of biotinylated lysozyme for 1 hour at room temperature followed by secondary labeling with SA-PE (1:250 dilution) for 12 minutes on ice and immunofluorescence detection. Using a similar approach, we also compared the binding of lysozyme and BSA (100, 500, 1000, and 2000 nM) to yeast cells displaying cyclic peptides of interest. BSA was biotinylated in a similar manner as previously described for lysozyme. The peptide expression level of each cell population was analyzed by labeling with a 1:100 dilution of an anti-HA antibody (Thermo Fisher Scientific) for 1 hour at room temperature followed by labeling with a 1:250 dilution of a donkey anti-rabbit 488 antibody (DAR488; Immunoreagents) for 12 minutes on ice and immunofluorescence detection. For each mutant, expression and secondary only (SA-PE and DAR488) labelings were performed for both the linear and cyclic populations. All labelings took place in 50 µL using 0.1% PBSA. Washes with 0.1% PBSA were performed after each incubation. The mean fluorescence of biotinylated lysozyme and biotinylated BSA binding was normalized by subtracting the mean fluorescence of the secondary SA-PE only labeling. Similarly, the HA expression mean fluorescence was normalized by subtracting the mean fluorescence of the secondary DAR488 only labeling. For each experimental sample, the background subtracted mean fluorescence values for lysozyme and BSA binding were normalized by dividing the background subtracted HA expression mean fluorescence value for that particular population. Error bars represent the standard error of the mean for three independent experiments.

*Screening a cyclic heptapeptide yeast display library against IL-17*

Cyclic peptides with affinity for IL-17 were discovered by performing magnetic selections against yeast displayed IL-17 followed by FACS selections against soluble, biotinylated IL-17. Initially, IL-17 was co-displayed on the yeast surface with SsoFe2, an iron oxide binding protein, using the pCT302-T2A-SSoFe2 plasmid as previously described[7]. To generate pCT302-T2A-SSoFe2-IL17, DNA corresponding to human IL-17 (Met1-Ala155, gene block 3) was amplified using primers Pf4 and Pr4 followed by insertion between the NheI and BamHI sites. For the magnetic selection, a negative selection against yeast cells expressing TOM22[7] and a positive selection against cells expressing IL-17 was performed. To begin, $7x10^8$ cells co-expressing IL-17 and SsoFe2 were resuspended in 10 mL of 1% PBS BSA (1% PBSA). Cells co-expressing TOM22 and SsoFe2 were also treated in a similar manner. The cells were magnetized by adding 2 mL of an iron oxide solution (4 mg/mL in water) followed by a 20-minute, rotating incubation at room temperature. Any unbound cells were removed. The magnetized cells were then washed 3X with 1% PBSA. Based on the $OD_{600}$ of the solution prior to the addition of the iron oxide and after the removal of the yeast bound to the iron oxide, $1x10^8$ magnetized cells were aliquoted for the magnetic selections. The aliquoted cells were blocked with 1% PBSA for 1 hour at room temp (10 mL). Subsequently, the TOM22 magnetized cells were washed 3X with 0.1% PBSA prior to the addition of $5x10^8$ cyclic heptapeptide library cells and

1x10$^9$ EBY100 cells in 10 mL of 0.1% PBSA. After an hour incubation, library cells that did not bind the magnetized TOM22 cells were recovered. The IL-17 magnetized cells were washed three times with 0.1% PBSA after blocking and then incubated for 1 hour with the cells recovered from the negative selection. Library cells bound to the magnetized IL-17 cells were isolated using a magnet. Magnetic IL-17 cells and any complexed library cells were washed 5 times with 0.1% PBSA prior to expansion in 20 mL of SDCAA (-TRP) media.

Using the library cells isolated from the magnetic screen, FACS selections were performed against soluble IL-17 in a similar manner as the FACS selections against lysozyme. Briefly, 1x10$^7$ cyclized, library cells were labeled with biotinylated IL-17A (R&D systems) for 1 hour on ice followed by secondary labeling using SA-PE (1:250 dilution) for 12 minutes on ice. The cells were labeled with 1 μM and 500 nM of biotinylated IL-17 for the first and second rounds of FACS, respectively. All labelings took place in 100 μL of 0.1% PBSA. Washes with 0.1% PBSA were performed after each incubation. DNA was extracted from the yeast cells isolated from the final IL-17 FACS round using a Zymoprep™ Yeast Plasmid Miniprep II kit following the manufacturer's recommendations. DNA encoding the isolated cyclic peptides was amplified from the extracted plasmid DNA using primers Pf5 and Pr5. Multiple rounds of PCR were performed to obtain the concentration of DNA required for sequencing. After the first round of PCR, amplified DNA was purified by phenol: chloroform extraction and concentrated using ethanol precipitation as previously described. For subsequent PCR rounds, the amplified DNA was purified using the BioBasic 9K Series Gel and PCR extraction kit. Partial Illumina adapters were added to the amplified DNA by Genewiz. The DNA was sequenced using Genewiz's Amplicon Ez protocol. Genewiz provided bioinformatic details about the recovered sequences, including how many times a read appeared. In total, 102,896 reads were obtained after performing Illumina sequencing on the final FACS population.

*In silico evaluation of the IL-17: cyclic peptide and IL-17:peptide:IL-17RA interactions*

Given the strong amphiphilic nature of the selected peptides, we used the hydropathy index (GRAVY) according to the Kyte-Doolittle scale to select specific peptides for *in silico* analysis from the Illumina sequencing results[8]. We chose peptides that were likely to be water soluble and exhibit low-nonspecific binding resulting from potential hydrophobic interactions (GRAVY < -0.5). The crystal structure of IL-17 (PDB ID: 4HSA)[9] was initially prepared using Schrödinger's Protein Preparation Wizard[10,11] to identify and correct missing atoms and/or side chains, remove salt ions and other small molecules, add explicit hydrogens, assign tautomeric states with EPIK, optimize the hydrogen-bonding networks, and minimize the protein energy using the OPLS force field[12,13]. The adjusted structure was subjected to a "druggability" study using SiteMap to identify putative binding sites capable of accommodating the cyclic heptapeptides[14–16]. The SiteMap algorithm by Schrödinger identifies binding pockets by locating spherical "site points" onto the protein surface; these points are clustered based on *(i)* their ability to form favorable protein-ligand interactions, *(ii)* solvent exposure, and *(iii)* hydrophobic/philic character. The regions on the protein surface that possess a sufficient number of site points and volume are scored using an S-score (likelihood of the protein's surface to be a binding pocket) and D-score (measure of the pocket's "druggability"). Sites with S-score and D-score values greater than or equal to 0.7 and 0.9, respectively, were selected as putative sites for ligand binding. In this work, given the topology of IL-17, the SiteMap analysis was performed with the '*detect shallow binding sites*' option selected, which adjusts amino acid atomic van der Waals radii to be more accommodating for peptide binding when locating potential binding pockets.

An ensemble, containing 17 cyclic peptides identified from the library screen, was designed using the molecular editor Avogadro[17,18], and equilibrated via atomistic molecular dynamics (MD) simulations in the GROMACS package[19–21] using the OPLS all-atom force field[22,23]. Every peptide was placed in a simulation box with periodic boundary conditions containing 800 water molecules (TIP3P water model)[24–26]. The system was minimized by running 10,000 steps of steepest gradient descent, heated to 300 K in an NVT ensemble for 250 ps (1 fs time steps), and equilibrated to 1 atm by running a 500-ps NPT simulation (2 fs time steps). The production run for every peptide was performed in the NPT ensemble at 300 K and 1 atm using the Nosé-Hoover thermostat[27–29] and Parrinello-Rahman barostat, respectively[30,31]. The leap-frog algorithm was used to integrate the equations of motion. All of the covalent bonds were constrained by means of the LINCS algorithm[32]. The short-range electrostatic and Lennard-Jones interactions were calculated within a cutoff of 1.0 nm and 1.2 nm, respectively, whereas the particle-mesh Ewald method was utilized to treat the long-range electrostatic interactions[33,34]. The atomic coordinates were saved every 2 ps, and the non-bonded interaction pair-list was updated every 5 fs using a cutoff of 1.4 nm.

The resulting peptide structures were then docked *in silico* against the selected putative binding sites on IL-17 using the docking software HADDOCK (High Ambiguity Driven Protein-Protein Docking) (v.2.1) [35–37]. The residues within IL-17's binding sites were defined as "active", whereas the residues surrounding the binding sites were defined as "passive". All variable amino acid positions on the peptide ligands were also denoted as 'active'. A GSG tripeptide segment located on the C-terminus of the peptide was defined as not involved in the interaction to account for the directionality of binding. When displayed on the surface of yeast, in fact, the peptide is connected to the Aga2p display system through its C-terminus. Docking in HADDOCK proceeded through a 3-stage protocol: *(i)* rigid, *(ii)* semi-flexible, and *(iii)* water-refined fully flexible docking. A total of 1000, 200, and 200 structures were calculated at each stage, respectively. Final structures were clustered using the cutoff of Cα RMSD < 7.5 Å as calculated in ProFit (Martin, A.C.R, http://www.bioinf.org.uk/software/profit/). The peptides in the identified clusters were ranked using FireDock[38,39] and Xscore[40,41]. The selected binding poses were finally refined via 100-ns atomistic molecular dynamics (MD) simulations using the GROMACS simulation package. The IL-17: peptide complexes were embedded in a cubic box of 9.7 nm side and periodic boundary and solvated with 30,000 TIP3P water molecules. The MD simulations were performed at 300 K and 1 atm using the Amber99SB force field. The MM/GBSA method was used for processing the refined peptide-IL-17 complexes and to estimate the free energy of binding ($\Delta G_B$)[42,43]. Finally, the IL-17:peptide:IL-17RA interactions were evaluated by docking IL-17RA (PDB: 3JVF)[44] on the MD-refined IL-17:peptide complexes; the active sites on IL-17RA were assigned based on the analysis of the IL-17:IL-17RA complex using the service PSIA ('Protein interfaces, surfaces, and assemblies) at the European Bioinformatics Institute (http://www.ebi.ac.uk/pdbe/prot_int/pistart.html)[45]. IL-17:peptide:IL-17RA complexes selected via FireDock-guided ranking[38,39] were refined via 200-ns atomistic molecular dynamics (MD) simulations, and processed by the MM/GBSA method to estimate $\Delta G_B$ values[42,43].

*Cloning of IL-17 cyclic peptides for yeast display*

Gene blocks 4 – 9 encoding the linear peptide sequences for cyclo-[*DSG*-RMRWLRGRR-*K*], cyclo-[*DSG*-RMIGQWWRR-*K*], cyclo-[*DSG*-RMNRLKFWF-*K*], cyclo-[*DSG*-RMSFFDIWR-*K*], cyclo-[*DSG*-RMYRFHRHG-*K*], and cyclo-[*DSG*-RMFGLLHRG-*K*], respectively, were amplified using primers Pf6 and Pr6. These gene blocks were digested and inserted between the EcoRI and XmaI cut sites of the pCT-NT-F2A-Sso7dhFc vector[4].

*Affinity and binding selectivity characterization of the IL-17 cyclic peptides*

The interaction of the yeast displayed IL-17 binding cyclic peptides with IL-17 and other putative, non-specific proteins was evaluated using flow cytometry analysis. Induced yeast cells displaying linear peptide sequences were cyclized using the DSG crosslinking protocol as previously described. EBY100 non-displaying yeast were also subjected to the DSG crosslinking protocol. The binding of biotinylated IL-17 at a concentration of 500 nM was studied for yeast displayed cyclo-[*DSG*-RMRWLRGRR-*K*], cyclo-[*DSG*-RMIGQWWRR-*K*], cyclo-[*DSG*-RMNRLKFWF-*K*], cyclo-[*DSG*-RMSFFDIWR-*K*], cyclo-[*DSG*-RMYRFHRHG-*K*], and cyclo-[*DSG*-RMFGLLHRG-*K*] as well as for DSG-treated EBY100 cells. The binding of biotinylated lysozyme (500 & 1000 nM), biotinylated BSA (500 & 1000 nM), GFP (500 & 1000 nM), and SA-PE (500 nM) was evaluated using immunofluorescence detection for yeast displayed cyclo-[*DSG*-RMRWLRGRR-*K*]. Binding of GFP and SA-PE at the described concentrations was also evaluated for the DSG-treated EBY100. All primary incubations using biotinylated protein took place for 30 minutes at room temperature. The binding of biotinylated protein was detected using a secondary SA-PE labeling (1:250 dilution, 12 minutes on ice in the dark). When evaluating the binding of GFP and SA-PE alone, no secondary detection agent was used as these proteins, by themselves, can provide a fluorescent signal when excited. When cells were only labeled with GFP and SA-PE, incubations took place in the dark for 30 minutes at room temperature. All labelings took place in a volume of 50 μL using 0.1% PBSA. Washes with 0.1% PBSA were performed after each incubation. The mean binding fluorescence of IL-17, lysozyme, and BSA was normalized by subtracting the background mean fluorescence of the secondary labeling agent and multiplying by a factor of 100. The mean fluorescence values of GFP and SA-PE binding were not background subtracted and only multiplied by a factor of 100. Error bars represent the standard error of the mean for three individual replicates.

*In vitro evaluation of cyclo-[DSG-MRWLRGRR-K] as a modulator*

Two approaches were used to study if cyclo-[*DSG*-MRWLRGRR-*K*] modulates the interaction between IL-17 and its receptor. In approach one, $2x10^6$ yeast cells displaying cyclo-[*DSG*-RMRWLRGRR-*K*] were incubated with 100 nM of biotinylated IL-17 in the presence or absence of 500 nM IL-17RA (R&D systems) overnight at 4°C in a total volume of 100 μL using 0.1% PBSA. The following day, the cells were washed and incubated with a 1:250 dilution of SA-PE for 12 minutes on ice (50 μL labeling in 0.1% PBSA). SA-PE binding was detected using flow cytometry. A secondary only labeling using SA-PE alone was also performed. The mean fluorescence of IL-17 binding was normalized by subtracting the mean fluorescence of the secondary only labeling control and multiplied by a factor of 100. A two-tailed, two sample unequal variance t-test was performed using individual data points from the flow cytometry analysis (n=390,763) comparing the mean fluorescence of the IL-17 binding of populations where IL-17RA was present or absent. Data points used in the t-test were not background subtracted. The binding of IL-17 to the yeast cells displaying cyclo-[*DSG*-RMRWLRGRR-*K*] was statistically different when the receptor was present ($p<0.001$).

The second approach differed by using soluble peptide and yeast displayed IL-17. The UNC High-Throughput Peptide Synthesis and Array facility synthesized a soluble form of cyclo-[*DSG*-MRWLRGRR-*K*][3]. This peptide lacked the N-terminal arginine residue present for the yeast displayed peptide sequences. To confer expression of IL-17 as a yeast surface fusion, gene block 3 was amplified using primers Pf4 and Pr4 and inserted between the NheI and BamHI sites of pCTCON to generate pCTCON-IL17[46]. $2x10^6$ yeast cells displaying IL-17 were incuabted overnight at 4°C with varying concentrations of soluble cyclo-[*DSG*-MRWLRGRR-*K*] (20 nM – 50 μM) and either 20 or 50 nM of IL-17RA. All incubations took place in 100 μL of 0.1% PBSA. A control was also performed for each concentration of IL-17RA where no peptide was present. The next morning the cells were incubated with a 1:100 dilution of biotinylated protein A (Thermo Fisher Scientific) for 30 minutes at room temperature (50 µL labeling in 0.1% PBSA). Thereafter, the cells were labeled with a 1:250 dilution of SA-PE for 12 minutes on ice (50 µL labeling in 0.1% PBSA), and subsequently analyzed by flow cytometry. A wash with 0.1% PBSA took place between all incubations. A labeling control was also performed where cells were incubated with biotinylated protein A followed by SA-PE in a similar fashion. The mean fluorescence of IL-17RA binding was normalized by subtracting the mean fluorescence of this control and multiplying by 100.

Additionally, a control experiment was performed where yeast cells displaying cyclo-[*DSG*-RMRWLRGRR-*K*] were incubated with IL-17RA. 2x$10^6$ yeast cells displaying cyclo-[*DSG*-RMRWLRGRR-*K*] were incuabted with 500 nM of IL-17RA overnight at 4ºC followed by labeling with biotinylated protein A (1:100 dilution) for 30 minutes at room temperature and SA-PE labeling for 12 minutes on ice in the dark (1:250 dilution). Non-displaying EBY100 yeast subjected to the DSG crosslinking procedure were also labeled in a similar manner. All labeling took place in 50 µL using 0.1% PBSA, and washes took place between each incubation with 0.1% PBSA. Binding was detected using flow cytometry. A labeling control was also performed where yeast cells displaying cyclo-[*DSG*-RMRWLRGRR-*K*] and DSG-treated EBY100 were incubated with biotinylated protein A (1:100 dilution) and SA-PE (1:250 dilution) following a similar procedure. The mean fluorescence of IL-17RA binding was normalized by subtracting the mean fluorescence of this control and multiplying by 100.

**Supplementary Figures and Tables**

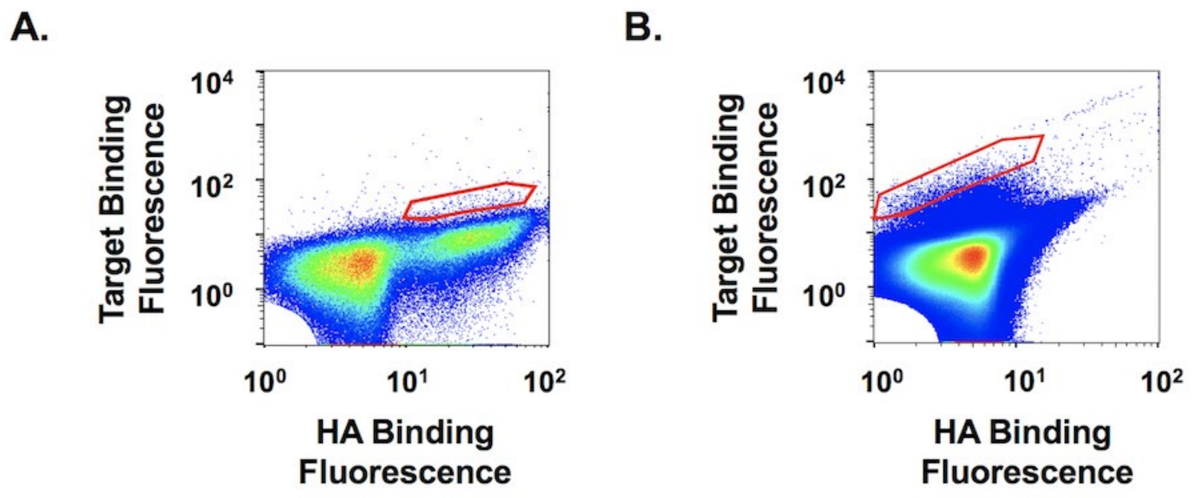


**Figure S1**. Comparison of **(A)** double labeled population for target binding and HA expression and **(B)** single labeled population for target binding only when performing fluorescence activated cell sorting (FACS) of yeast displayed cyclic peptide libraries.

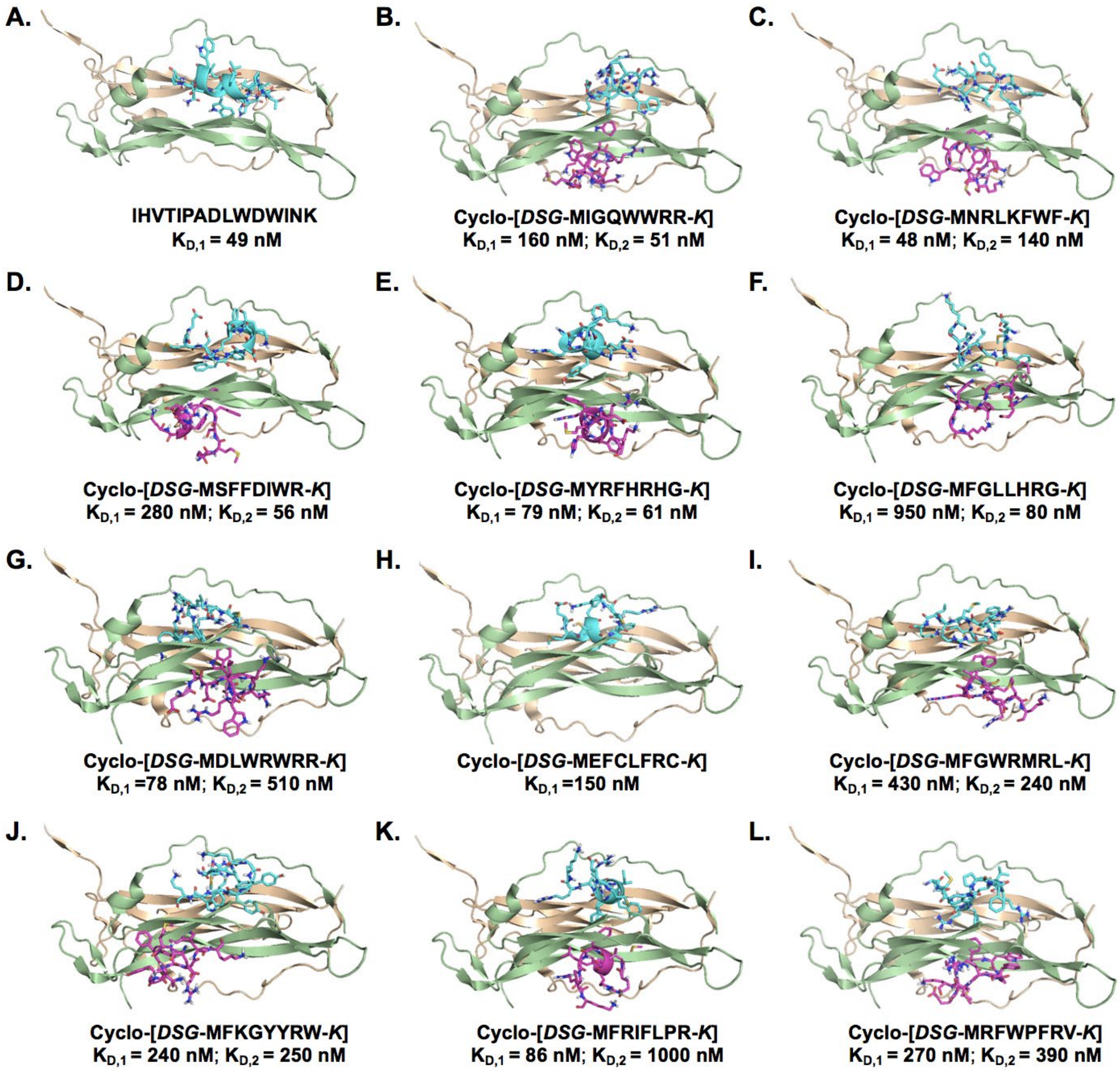


**Figure S2**. *In silico* IL-17: cyclic peptide complexes obtained by molecular docking and refined by molecular dynamic simulations. The monomers of IL-17 are denoted in green and brown while the respective cyclic peptide is modeled in cyan or magenta. The positive control, linear peptide, IHVTIPADLWDWINK, is modeled in **(A)**.

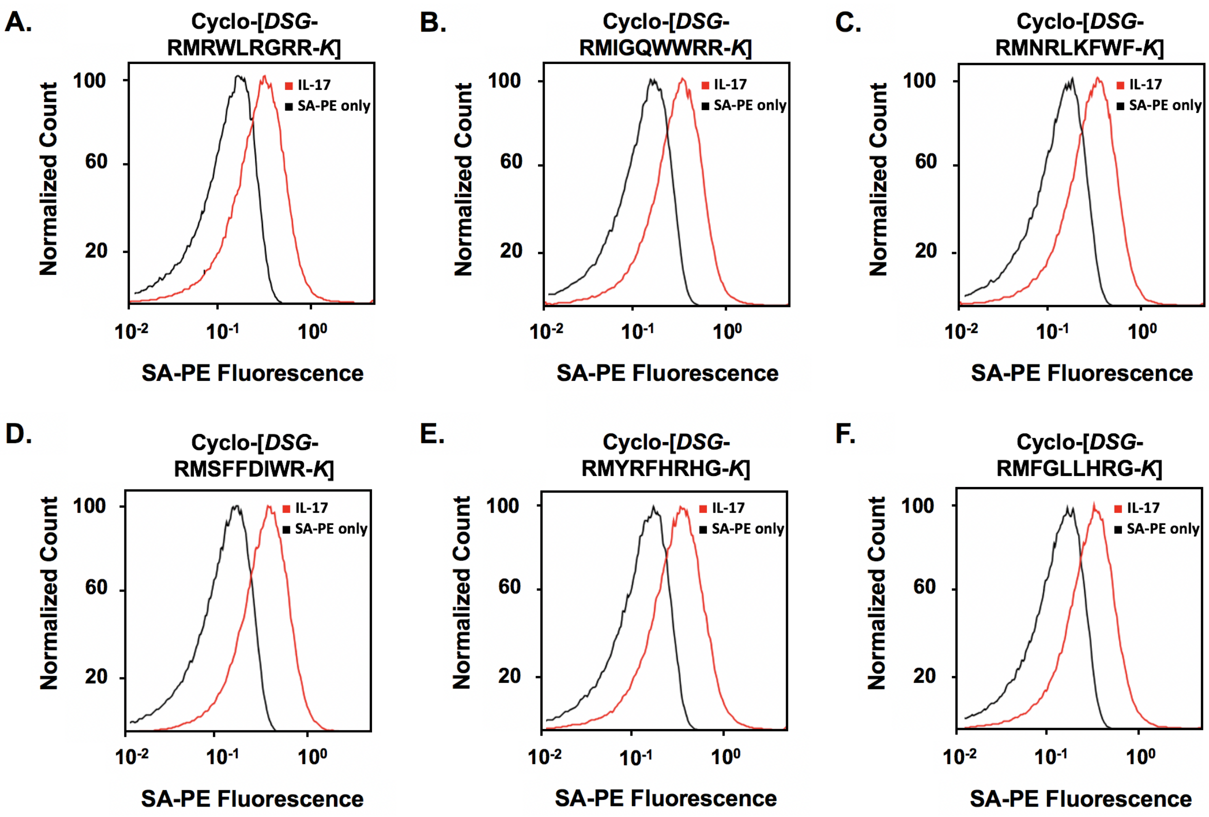


**Figure S3**. Binding of IL-17 (500 nM) to yeast displayed cyclic peptides (red) detected using SA-PE. The binding of SA-PE alone was also considered (black). Cyclic peptides considered include: **(A)** cyclo-[*DSG*-RMRWLRGRR-*K*], **(B)** cyclo-[*DSG*-RMIGQWWRR-*K*], **(C)** cyclo-[*DSG*-RMNRLKFWF-*K*], **(D)** cyclo-[*DSG*-RMSFFDIWR-*K*], **(E)** cyclo-[*DSG*-RMYRFHRHG-*K*], and **(F)** cyclo-[*DSG*-RMFGLLHRG-*K*].

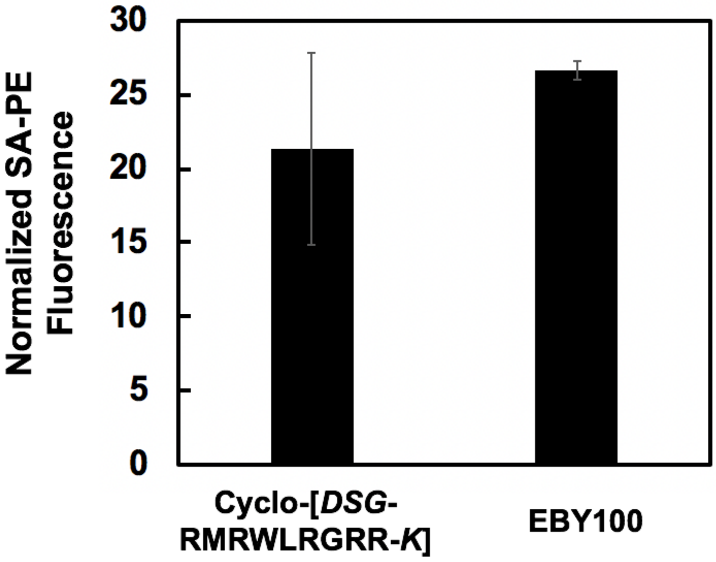


**Figure S4**. Cyclo-[*DSG*-RMRWLRGRR-*K*] does not interact with IL-17RA. Yeast cells displaying cyclo-[*DSG*-RMRWLRGRR-*K*] and DSG-treated EBY100 yeast were incubated with 500 nM of IL-17RA followed by detection with biotinylated protein A and SA-PE. The mean fluorescence of IL-17RA binding was normalized by the mean fluorescence of biotinylated protein A and SA-PE binding to each population. Error bars represent the standard error of the mean for three individual repeats.

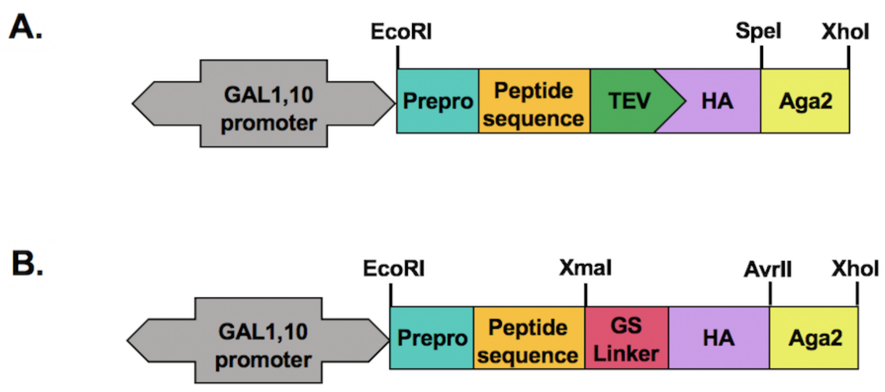


**Figure S5**. Plasmid design for display of **(A)** IgG binding peptides and **(B)** variants from the heptapeptide yeast display library.

**Table S1**. Sequences of IL-17 binding cyclic peptides analyzed in a molecular docking study with the IL-17 interface. Mutagenized positions are denoted with an X. Numbers in parentheses represent the number of times a sequence appeared during Illumina Next Generation Sequencing.

| | M | $X_1$ | $X_2$ | $X_3$ | $X_4$ | $X_5$ | $X_6$ | $X_7$ | K |
|---|---|---|---|---|---|---|---|---|---|
| **IL17.1 (4191)** | | R | W | L | R | G | R | R | |
| **IL17.2 (1197)** | | F | G | L | L | H | R | G | |
| **IL17.3 (1082)** | | D | L | W | R | W | R | R | |
| **IL17.4 (747)** | | S | F | R | F | W | R | L | |
| **IL17.5 (498)** | | S | F | F | D | I | W | R | |
| **IL17.6 (428)** | | I | G | Q | W | W | R | R | |
| **IL17.7 (318)** | | R | F | W | P | F | R | V | |
| **IL17.8 (307)** | | F | L | F | R | L | G | R | |
| **IL17.9 (285)** | | E | F | C | L | F | R | C | |
| **IL17.10 (275)** | | Y | R | F | H | R | H | G | |
| **IL17.11 (251)** | | F | R | I | F | L | P | R | |
| **IL17.12 (229)** | | C | L | L | W | C | R | R | |
| **IL17.13 (228)** | | F | K | G | Y | Y | R | W | |
| **IL17.14 (219)** | | N | R | L | K | F | W | F | |
| **IL17.15 (145)** | | K | H | R | P | F | R | Y | |
| **IL17.16 (141)** | | F | G | W | R | M | R | L | |
| **IL17.17 (119)** | | P | F | G | H | W | R | R | |

**Table S2**. Values of binding energy ($\Delta G_B$) and calculated affinity ($K_D$) of the IL-17: peptide complexes derived from docking and molecular dynamic simulations. IHVTIPADLWDWINK and cyclo-[*DSG*-MGSGGGGSG-K] were used as positive and negative controls, respectively.

| Sequence | Site 1 $\Delta G_B$ (kcal/mol) | Site 1 $K_D$ (nM) | Site 2 $\Delta G_B$ (kcal/mol) | Site 2 $K_D$ (nM) |
|---|---|---|---|---|
| IHVTIPADLWDWINK | -9.97 | 49 | | |
| Cyclo-[*DSG*-MGSGGGGSG-*K*] | -5.57 | 81000 | | |
| Cyclo-[*DSG*-MRWLRGRR-*K*] | -10.14 | 36 | -9.59 | 91 |
| Cyclo-[*DSG*-MIGQWWRR-*K*] | -9.26 | 160 | -9.93 | 51 |
| Cyclo-[*DSG*-MNRLKFWF-*K*] | -9.97 | 48 | -9.34 | 140 |
| Cyclo-[*DSG*-MSFFDIWR-*K*] | -8.93 | 280 | -9.88 | 56 |
| Cyclo-[*DSG*-MYRFHRHG-*K*] | -9.67 | 79 | -9.83 | 61 |
| Cyclo-[*DSG*-MFGLLHRG-*K*] | -8.20 | 950 | -9.67 | 80 |
| Cyclo-[*DSG*-MDLWRWRR-*K*] | -9.68 | 78 | -8.57 | 510 |
| Cyclo-[*DSG*-MEFCLFRC-*K*] | -9.29 | 150 | | |
| Cyclo-[*DSG*-MFGWRMRL-*K*] | -8.67 | 430 | -9.02 | 240 |
| Cyclo-[*DSG*-MFKGYYRW-*K*] | -9.02 | 240 | -9.00 | 250 |
| Cyclo-[*DSG*-MFRIFLPR-*K*] | -9.63 | 86 | -8.18 | 1000 |
| Cyclo-[*DSG*-MRFWPFRV-*K*] | -8.96 | 270 | -8.74 | 390 |

**Table S3**. List of gene block fragments

| | |
|---|---|
| Gene Block 1 | GAATTCATGAAGGTTTTGATTGTCTTGTTGGCTATCTTCGCTGCTTTGCCA<br>TTGGCCTTAGCTCAACCAGTAATTTCTACTACCGTCGGTTCCGCTGCAGA<br>AGGCTCTTTGGACAAGAGAGCTAGCGGACAATTACTATTTACAATTACAA<br>TGAGTCATAGAGCAGGTCTGGAGAAGGGCAGCGGCGGCAGCGGATCCGA<br>GAACCTGTACTTCCAAGGGTATCCGTACGACGTTCCAGACTACGCTACTA<br>GTCAGGAACTGACAACTATATGCGAGCAAATCCCCTCACCAACTTTAGA<br>ATCGACGCCGTACTCTTTGTCAACGACTACTATTTGGCCAACGGGAAGG<br>CAATGCAAGGAGTTTTTGAATATTACAAATCAGTAACGTTTGTCAGTAAT<br>TGCGGTTCTCACCCCTCAACAACTAGCAAAGGCAGCCCCATAAACACAC<br>AGTATGTTTTTTAATGACTCGAG |
| Gene Block 2 | GAATTCATGAAGGTTTTGATTGTCTTGTTGGCTATCTTCGCTGCTTTGCCA<br>TTGGCCTTAGCTCAACCAGTAATTTCTACTACCGTCGGTTCCGCTGCAGA<br>AGGCTCTTTGGACAAGAGAATGNNKNNKNNKNNKNNKNNKNNKAAGCC<br>CGGGGGTGGTGGTGGTTCTGGAGGAGGCTCTGGTG |
| Gene Block 3 | ATGACGCCTGGGAAAACGAGCTTGGTCTCACTACTATTGCTTCTGAGCCT<br>TGAGGCGATAGTTAAGGCGGGTATCACAATACCTAGAAACCCAGGCTGC<br>CCGAACAGTGAAGACAAAAACTTCCCGAGAACTGTGATGGTCAACCTAA<br>ACATACACAATAGGAATACAAACACTAATCCAAAGAGGAGCAGTGATTA<br>CTACAATAGATCAACCTCTCCTTGGAACTTACATCGTAATGAGGACCCTG<br>AGAGGTATCCTTCAGTCATCTGGGAGGCTAAGTGTAGACATTTGGGCTGC<br>ATCAACGCTGACGGTAACGTCGATTATCACATGAACTCCGTGCCTATTCA<br>ACAGGAAATTCTGGTATTGAGGCGTGAGCCTCCTCATTGTCCCAATAGTT<br>TCAGGCTTGAGAAGATCCTTGTTTCAGTTGGCTGCACTTGCGTCACTCCA<br>ATCGTACACCACGTTGCA |
| Gene Block 4 | GAATTCATGAAGGTTTTGATTGTCTTGTTGGCTATCTTCGCTGCTTTGCCA<br>TTGGCCTTAGCTCAACCAGTAATTTCTACTACCGTCGGTTCCGCTGCAGA<br>AGGCTCTTTGGACAAGAGAATGCGTTGGTTAAGGGGAAGAAGGAAACCC<br>GGGGGTGGTGGTGGTTCT |
| Gene Block 5 | GAATTCATGAAGGTTTTGATTGTCTTGTTGGCTATCTTCGCTGCTTTGCCA<br>TTGGCCTTAGCTCAACCAGTAATTTCTACTACCGTCGGTTCCGCTGCAGA<br>AGGCTCTTTGGACAAGAGAATGATAGGCCAATGGTGGAGAAGAAAGCCC<br>GGGGGTGGTGGTGGTTCT |
| Gene Block 6 | GAATTCATGAAGGTTTTGATTGTCTTGTTGGCTATCTTCGCTGCTTTGCCA<br>TTGGCCTTAGCTCAACCAGTAATTTCTACTACCGTCGGTTCCGCTGCAGA<br>AGGCTCTTTGGACAAGAGAATGAACCGTCTTAAATTTTGGTTTAAACCCG<br>GGGGTGGTGGTGGTTCT |
| Gene Block 7 | GAATTCATGAAGGTTTTGATTGTCTTGTTGGCTATCTTCGCTGCTTTGCCA<br>TTGGCCTTAGCTCAACCAGTAATTTCTACTACCGTCGGTTCCGCTGCAGA<br>AGGCTCTTTGGACAAGAGAATGTCCTTCTTCGACATATGGCGTAAACCCG<br>GGGGTGGTGGTGGTTCT |
| Gene Block 8 | GAATTCATGAAGGTTTTGATTGTCTTGTTGGCTATCTTCGCTGCTTTGCCA<br>TTGGCCTTAGCTCAACCAGTAATTTCTACTACCGTCGGTTCCGCTGCAGA |

| | |
|---|---|
| | AGGCTCTTTGGACAAGAGAATGTACCGTTTCCATAGGCATGGAAAGCCC GGGGGTGGTGGTGGTTCT |
| Gene block 9 | GAATTCATGAAGGTTTTGATTGTCTTGTTGGCTATCTTCGCTGCTTTGCCA TTGGCCTTAGCTCAACCAGTAATTTCTACTACCGTCGGTTCCGCTGCAGA AGGCTCTTTGGACAAGAGAATGTTTGGCTTGTTGCACCGTGGCAAGCCCG GGGGTGGTGGTGGTTCT |

**Table S4**. List of oligonucleotide primers

| Primer | Sequence |
|---|---|
| Pf1 | CTGGACGAATTCATGAAGGTTTTGATTGTCTT |
| Pr1 | CTGGACCTCGAGTCATTAAAAAACATACTGT |
| Pf2 | GCAGAAGGCTCTTTGGACAA |
| Pr2 | ATACCCTTGGAAGTACAGGTT |
| Pf3 | GTATTACTTCTTATTCAAATGTAATAAAAGATCGAATTCATGAAGGTTTTG ATTGTCTTGTTG |
| Pr3 | CACCAGAGCCTCCTCCAGA |
| Pf4 | CATAGCGCTAGCATGACGCCTGGGAAAACGAG |
| Pr4 | CACTGCGGATCCTGCAACGTGGTGTACGATTGG |
| Pf5 | GAATTCATGAAGGTTTTGATTGTCT |
| Pr5 | AGCGTAGTCTGGAACGTCGT |
| Pf6 | GCACGAGAATTCATGAAGGTTTTGATTGTC |
| Pr6 | GCACGAAGAACCACCACCACCCC |

**Table S5**. Oligonucleotide DNA fragments

| Oligo | Sequence |
|---|---|
| Oligo 1 | GCAGAAGGCTCTTTGGACAAGAGAATGTGGTTTCCGCATTACAAGGAGAAC CTGTACTTCCAAGGGTAT |
| Oligo 2 | GCAGAAGGCTCTTTGGACAAGAGAATGCACGGTTTTAGAGGCAAGGAGAAC CTGTACTTCCAAGGGTAT |
| Oligo 3 | GCAGAAGGCTCTTTGGACAAGAGAATGTGGTTTAGGCACTACAAGGAGAAC CTGTACTTCCAAGGGTAT |

(17) Hanwell, M. D.; Curtis, D. E.; Lonie, D. C.; Vandermeerschd, T.; Zurek, E.; Hutchison, G. R. Avogadro: An Advanced Semantic Chemical Editor, Visualization, and Analysis Platform. *J.*

*Cheminform.* 2012, *4* (8), 17.

(18) Avogadro: An Open-Source Molecular Builder and Visualization Tool. Version 1.2.0 Http://Avogadro.Cc/.

(35) Dominguez, C.; Boelens, R.; Bonvin, A. M. J. J. HADDOCK: A Protein-Protein Docking Approach Based on Biochemical or Biophysical Information. *J. Am. Chem. Soc.* 2003, *125* (7),

1731–1737.
(36) De Vries, S. J.; Van Dijk, M.; Bonvin, A. M. J. J. The HADDOCK Web Server for Data-Driven Biomolecular Docking. *Nat. Protoc.* 2010, *5* (5), 883–897.
(37) Van Zundert, G. C. P.; Rodrigues, J. P. G. L. M.; Trellet, M.; Schmitz, C.; Kastritis, P. L.; Karaca, E.; Melquiond, A. S. J.; Van Dijk, M.; De Vries, S. J.; Bonvin, A. M. J. J. The HADDOCK2.2 Web Server: User-Friendly Integrative Modeling of Biomolecular Complexes. *J. Mol. Biol.* 2016, *428* (4), 720–725.
(38) Mashiach, E.; Schneidman-Duhovny, D.; Andrusier, N.; Nussinov, R.; Wolfson, H. J. FireDock: A Web Server for Fast Interaction Refinement in Molecular Docking. *Nucleic Acids Res.* 2008, *36* (Web Server issue), W229-32.
(39) Andrusier, N.; Nussinov, R.; Wolfson, H. J. FireDock: Fast Interaction Refinement in Molecular Docking. *Proteins Struct. Funct. Genet.* 2007, *69* (1), 139–159.
(40) Wang, R.; Lai, L.; Wang, S. Further Development and Validation of Empirical Scoring Functions for Structure-Based Binding Affinity Prediction. *J. Comput. Aided. Mol. Des.* 2002, *16* (1), 11–26.
(41) Wang, R.; Lu, Y.; Wang, S. Comparative Evaluation of 11 Scoring Functions for Molecular Docking. *J. Med. Chem.* 2003, *46* (12), 2287–2303.
(42) Hou, T.; Wang, J.; Li, Y.; Wang, W. Assessing the Performance of the MM/PBSA and MM/GBSA Methods. 1. The Accuracy of Binding Free Energy Calculations Based on Molecular Dynamics Simulations. *J. Chem. Inf. Model.* 2011, *51* (1), 69–82.
(43) Genheden, S.; Ryde, U. The MM/PBSA and MM/GBSA Methods to Estimate Ligand-Binding Affinities. *Expert Opin. Drug Discov.* 2015, *10* (5), 449–461.
(44) Ely, L. K.; Fischer, S.; Garcia, K. C. Structural Basis of Receptor Sharing by Interleukin 17 Cytokines. *Nat. Immunol.* 2009, *10* (12), 1245–1251.
(45) Krissinel, E.; Henrick, K. Inference of Macromolecular Assemblies from Crystalline State. *J. Mol. Biol.* 2007, *372* (3), 774–797.
(46) Gera, N.; Hill, A. B.; White, D. P.; Carbonell, R. G.; Rao, B. M. Design of PH Sensitive Binding Proteins from the Hyperthermophilic Sso7d Scaffold. *PLoS One* 2012, *7* (11), e48928.
